# AETA peptide contributes to Alzheimer’s disease signature of synapse dysfunction

**DOI:** 10.1101/2025.08.22.671719

**Authors:** Jade Dunot, Carine Gandin, Marin Truchi, Giulia Pirro, Sébastien Moreno, Agathe Launay, Benjamin Azoulay, Hugo Landra, Sandy Ma Yishan, Luc Buée, Kevin Lebrigand, Paula A. Pousinha, David Blum, Bernard Mari, Ingrid Bethus, Michael Willem, Hélène Marie

## Abstract

Alzheimer’s disease (AD), the leading cause of dementia, is characterized by early synaptic dysfunction that precedes overt cognitive decline. While amyloid-β and Tau remain central to AD pathogenesis, molecular triggers of synapse weakening remain unclear. Here, we investigated AETA, a novel brain-secreted peptide derived from amyloid precursor protein (APP), as a potential mediator of synapse dysfunction in AD. We previously identified AETA as a unique modulator of NMDA receptor activity in the healthy brain; however, its role in AD etiology was yet to be explored. Post-mortem analyses of human hippocampal and prefrontal cortex tissues revealed significantly elevated AETA levels in AD patients, particularly in females. To further explore the contribution of AETA to AD synaptic pathology, we analyzed a new mouse model, the AETA-m mouse, exhibiting chronically increased brain AETA expression. Hippocampi of female AETA-m mice display an increase in the number of astrocyte and microglia, but no overt neuroinflammation. RNA sequencing of female AETA-m hippocampi revealed alterations in synaptic gene expression that closely paralleled those observed in vulnerable human AD brain regions, most notably in the hippocampus. These two phenotypes were absent in males. Functionally, hippocampal neurons from AETA-m mice displayed impaired NMDA receptor signaling, dendritic spine loss, and memory deficits especially in females, mirroring early AD-associated synaptic dysfunction. Together, these findings identify AETA as a novel key contributor of synaptic vulnerability in AD and associated memory processing, especially in females. Targeting AETA signaling may therefore offer new therapeutic avenues for preventing or mitigating synaptic and cognitive decline in AD.

## Introduction

Alzheimer’s disease (AD), the most prevalent form of dementia, is clinically characterized by progressive memory decline [3]. Post-mortem examination of AD brains consistently reveals hallmark pathological features: extracellular amyloid-β (Aβ) plaques and intracellular neurofibrillary tangles composed of hyperphosphorylated Tau proteins [37]. While Aβ and Tau remain central to AD pathogenesis, growing evidence suggests that they act synergistically within a broader landscape of molecular disruptions, particularly during the early, preclinical ‘cellular phase’ of AD [10].

Aβ peptides originate from the proteolytic cleavage of the amyloid precursor protein (APP) through the amyloidogenic processing pathway. In contrast, non-amyloidogenic processing does not yield Aβ. A third APP cleavage route, the η-secretase pathway, was identified more recently [51] (Figure 1a). This pathway involves cleavage by η-secretase at the N-terminal extracellular domain of APP, generating soluble APPη and a membrane-bound C-terminal fragment (CTFη). Subsequent cleavage of CTFη by α- or β-secretases produces distinct Aη peptides (Aη-α and Aη-β), collectively termed AETA.

**Fig. 1.**
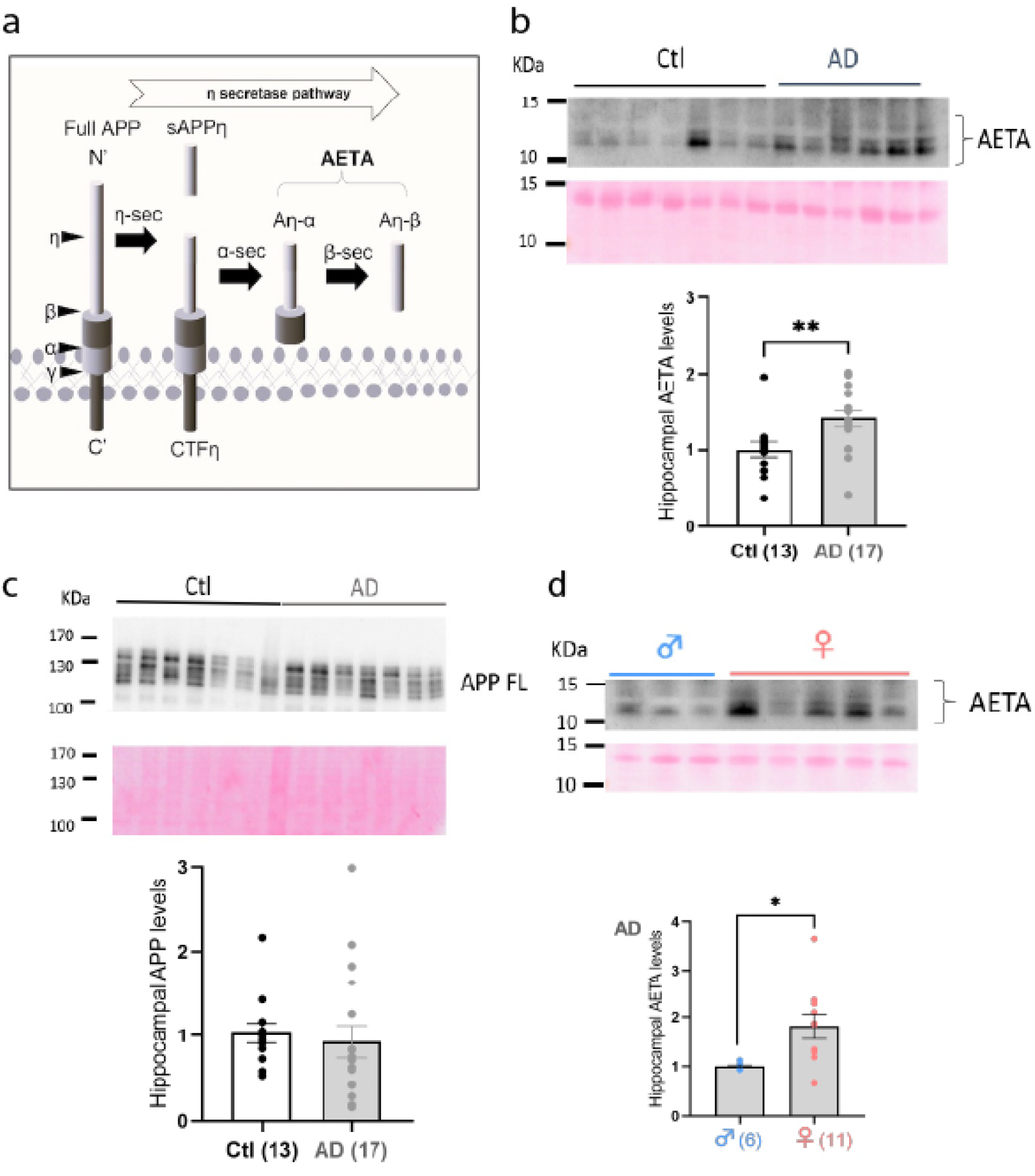
AETA levels are increased in hippocampus of AD patients. **(a)** Diagram depicting η-secretase processing and production of AETA peptides. **(b)** Immunoblot (2D8 antibody, which only recognizes the long α form of AETA, and ponceau staining for normalization) example and quantification of AETA levels (relative levels normalized to age-matched control (Ctl) subjects) in hippocampi of AD patient vs control subjects**. (c)** Immunoblot (Y188 antibody and ponceau staining for normalization) example and quantification of APP levels in the hippocampus (relative levels normalized to age-matched control (Ctl) subjects). **(d)** Immunoblot (2D8 antibody and ponceau staining for normalization) example and quantification of AETA levels in hippocampi of male vs female AD patients. Full blots used for analysis are provided in Supplemental material. N= number of human samples; each sample was tested in triplicate and presented as mean. Error bars represent s.e.m.; * p < 0.05; ** p < 0.01. Statistics: Mann-Whitney (b-c) and welch test (d). See full statistics in Supplemental Table S8.

We previously demonstrated that AETA impairs synaptic function by inhibiting long-term potentiation (LTP) at CA3-CA1 synapses and suppressing hippocampal calcium dynamics *in vivo* [28, 51]. More recently, we elucidated the molecular mechanism of the acute action of AETA at excitatory synapses [11]. AETA partially inhibits the ionotropic activity of NMDA receptors (NMDARs), by competing with the co-agonists glycine and D-serine. Concurrently, AETA promotes a non-canonical, ion flux-independent signaling mode of NMDARs, ultimately leading to a rapid reduction of synaptic strength and dendritic spine retraction. This AETA-dependent regulation of NMDARs is necessary in the healthy brain as, when AETA production is prevented, synapses cannot weaken when submitted to a long-term depression protocol [11].

Synaptic dysfunction and dendritic spine loss in the hippocampus are among the earliest and most predictive pathological changes in AD, tightly associated with cognitive decline and linked mechanistically to NMDAR signaling abnormalities [17, 22, 24, 40, 41, 45]. Several culprits have been previously identified as contributors to this synapse dysfunction including Aβ, Tau, mitochondrial dysfunction and microglia and astrocytes [17, 45]. However, the precise molecular triggers of this dysfunction remain to be fully elucidated, hindering targeted therapeutic development for treatment of AD. Notably, how AETA contributes to AD is currently unknown. Here we show that AETA increases in human AD patient brains. Mimicking this elevation in the rodent brain leads to human AD-like molecular and functional signatures of synapse dysfunction associated with hippocampus-dependent memory decay, particularly in female mice. These findings position AETA as a critical mediator of synaptic pathology in AD and highlight its potential as a novel therapeutic target.

## Materials and Methods

### Human brain samples

All experiments were performed in accordance with relevant guidelines and regulations. The case–control study of post-mortem human brain was approved by ministère de l’enseignement supérieur, de la recherche et de l’innovation (CODECOH N° DC-2022-5317). Human hippocampi were obtained from The Netherlands Brain Bank (NBB), Netherlands Institute for Neuroscience, Amsterdam, Netherlands (open access: www.brainbank.nl). Human prefrontal cortex samples were obtained from the Neuro-CEB Brain bank, Hôpital de la Pitié-Salpêtrière, Paris, France (open access: www.neuroceb.org). All Material was collected from donors for or from whom a written informed consent for a brain autopsy and the use of the material and clinical information for research purposes had been obtained by the NBB and Neuro-CEB. Patients had ante-mortem evidence of clinical dementia, whereas controls did not. Controls were selected by matching for age and sex (see Supplemental Tables S1 and S2 for details).

### Mice

All experiments were performed on 6-10 months old mice, unless specified in figure legend. Both sexes were used throughout the study and indicated for each experiment in legends and statistical analysis tables. All experiments and protocols on mice were performed in accordance with specific European laws described in the European Communities’ Council Directive 2010/63/EU. Protocols used in this study were approved by the committee for the Care and Use of Laboratory Animal in the framework of project authorizations APAFIS#6856-2016091610462338 and APAFIS#37493-2022050311352580s. Mice had ad libitum access to tap water and standard chow. Mice were maintained under constant environmental conditions (12h:12 h light/dark cycle, 23 ± 2°C and humidity of 55%). They were housed in groups of 5 or 6 animals by sex in standard mouse cages (542 cm^2^) under Specific Pathogen Free (SPF) status. Sex used is reported for each experiment.

### Generation and maintenance of AETA-m mouse line

Generation of AETA-m mouse line was briefly described previously [11]. Human AETA (the long alpha form) was first cloned into the multiple cloning site of pSecTag2 A plasmid downstream of the Ig k-chain leader sequence for secretion (Invitrogen Life Technologies, France) and then cloned into cDNA inserted in the mouse *thy-1* gene cassette at the *Xho*I site of the pTSC21k plasmid [2, 29]. This expression cassette was then linearized and injected into male pronuclei of fertilized zygotes of C57Bl6/NCrl donor mice. Embryos were transferred into Crl:CD1(ICR) pseudopregnant recipient female mice (ca. 20 embryos per recipient). The recipient mice were mated with sterile males (vasectomized) Crl:CD1(ICR). The AETA-m mouse line was chosen amongst four founder lines as it displayed moderate expression of recombinant AETA in the hippocampus and cortex (data not shown). Mice were generated under the license 24-9168.11-9/2012-5. All animal experiments were performed in accordance with the European Communities Council Directive (86/609/EEC) and were approved by the local ethics committee (Government of Saxony, Germany). The line was backcrossed regularly (at least 8 generations prior to use of mice for experiments) on C57/BL6J background strain (Charles River, France). Mouse genotype was confirmed by genomic DNA extraction from tissue and PCR as follows: primers sequences: CGCGCCATGATTAGTGAA and GACCTCTGCAGAGGAAGGA; PCR protocol: 5 min 95°C, 10 cycles of 94°C-30s/62°C-30s/72°C-30s, 24 cycles of 94°C-30s/52°C-30s/72°C-30s, 7 min 72°C. PCR samples were run on 2% agarose gel containing ethidium bromide and visualized by UV light.

### Expression of pSeg-Tag 2A-human AETA cDNA in CHO cells

CHO cells were either untransfected or transiently transfected with the pSeg-Tag 2A-human AETA cDNA with lipofectamine. Conditioned OPTIMEM medium from both plates was harvested after 48 hrs of expression. The collected media (from control untransfected or transfected CHO cells) were split in half and were directly loaded on two 12% Tris-Glycine gels for immunoblotting with 2E8 or 2E9 as detailed in immunoblotting of APP peptides section below.

### Collection of cerebrospinal fluid (CSF) from cisterna magna of mice

Collection of cerebrospinal fluid were performed using a stereotaxic frame (Kopf Instruments) under general anesthesia with xylazine and ketamine (10mg/kg and 150 mg/kg, respectively). The head was set with a 35° angle down. To expose the cisterna magna, a sagittal incision of the skin, inferior to the skull, was made and the subcutaneous tissue and neck muscles were separated through the midline. Two warming steps were used to prepare a capillary glass (step 1: 65.1°C; step 2: 52,7°C). The large end of the capillary glass was attached to a 1ml syringe to apply suction. This sharp end of the capillary glass was used to puncture the cisterna magna. The CSF was then transferred into a tube and snap frozen in liquid nitrogen.

### Immunoblotting of APP peptides in mouse and human tissues

Mouse and human brain homogenates for analysis of APP processing products (AETA and APP-FL (full-length) for human brain; AETA, APP-FL, sAPPα/β and CTF-η for mouse brains) were essentially prepared as described previously [51]. In brief, DEA lysates (0,2% Diethylamine in 50 mM NaCl, pH 10) and RIPA lysates (20 mM Tris-HCl pH 7.4, 150 mM NaCl, 1 mM Na2EDTA, 1% NP-40, 0,5 % sodium deoxycholate, 0,05 % Triton X-100) with protease inhibitors (Sigma-Aldrich, P8340) were prepared from brain samples using the Precellys system (Bertin) for homogenisation followed by ultracentrifugation. For the DEA and RIPA samples we used the Bradford (Biorad) to measure protein concentration, which was adjusted equally to all samples before immunoblotting. AETA (human and mouse) and sAPPα/β (mouse) and GAPDH (human) were quantified in DEA fraction and CTF-η (mouse) and full-length APP (human and mouse) were quantified in RIPA fraction. For detection by immunoblotting, proteins were separated on 8% Tris-Glycine gels or alternatively on Tris-Tricine (10-20%, Thermo Fisher Scientific) gels, transferred to nitrocellulose membranes (0.22 μm, GE Healthcare) which were boiled for 5 min in phosphate buffer saline (PBS) and subsequently incubated with the blocking solution containing 0.2% I-Block (Thermo Fisher Scientific) and 0.1% Tween 20 (Merck) in PBS for 1 hour, followed by overnight incubation with antibody in the blocking solution. Antibody detection was performed using the corresponding anti-rat/mouse/rabbit-IgG-HRP conjugated secondary antibody (Thermo Fisher Scientific) and chemiluminescence detection reagent ECL (Thermo Fisher Scientific). Antibodies used for immunoblotting were: anti-GAPDH (#G9545, Sigma-Aldrich, rabbit IgG, 1/1000 dilution), M3.2 (Biolegend, #805701; mouse IgG, 1/5000 dilution) for detection of mouse AETA and CTF-η, 2D8 (Sigma-Aldrich, #MABN2273, rat IgG; 2 μg/mL) or 2E9 (Sigma-Aldrich, #MABN2295, rat IgG, 2 μg/mL) for detection of human AETA, Y188 (Abcam, ab32136; rabbit IgG, 1/1000 dilution) for detection of human and mouse APP-FL, and 22C11 (Merck, #MAB348; mouse IgG, 1/5000 dilution) for detection of mouse sAPPα/β. For the loading control, when necessary, we used an antibody specific to β-actin (Sigma-Aldrich, #A5316, mouse IgG, 1/5000 dilution) or ponceau staining (as stated in results description, see also Supplemental file with full blots). Ponceau staining was generally chosen for normalization as limitations in housekeeping genes have been previously evidenced when comparing normal and diseased tissues [12]. For quantification, peptide levels were normalized to WT average for each immunoblot. For quantification of recombinant human AETA in hippocampi of AETA-m mice, the standard curve was made using the following synthetic peptide (Peptide Specialty Laboratories; PSL GmbH; Heidelberg, Germany): MISEPRISYGNDALMPSLTETKTTVELLPVNGEFSLDDLQPWHSFGADSVPANTENEVEPVDARPAADRGLTTRPGSGLTNIKTEEISEVKMDAEFRHDSGYEVHHQK. The peptide was dissolved in dimethyl sulfoxide (DMSO) at 100 μM and placed at −80°C for long term storage and diluted to final concentration on day of experiment.

### Immunohistochemistry

For Cresyl violet staining, brains were quickly removed and maintained in 4% paraformaldehyde (PFA) for 24h, then washed in PBS. Brain sections (40µm) were sliced with a vibratome (Leica VT 1000S). Cresyl violet staining to label Nils bodies was performed on slices just after slicing procedure as follows. Slices were mounted on superfrost slides (VWR) and allowed to dry. Dried slices were stained for 3 min in cresyl violet solution (5g cresyl violet/3ml acetic acid/ 1L ddH_2_O, filtered), then well rinsed in ddH_2_O. Slices were further washed in successive baths 70, 90 and 100% Ethanol for 1 min each. Slices were dried again at room temperature and mounted in Entellan (Sigma-Aldrich). Pictures were taken under Olympus BX41 microscope at 10X magnification. Cell body layer thickness was calculated manually with Image J software with the experimenter blinded to mouse genotype.

For 2E9, GFAP and IBA-1 immunohistochemistry on hippocampal slices, mice were submitted to an intracardiac perfusion with saline (0.9% NaCl) to remove blood, and then with PFA (4%) to fix tissues. Brains were removed and maintained in PFA for 24h, then washed in phosphate buffer saline (PBS). Brain sections (30µm) were sliced with a vibratome (Leica VT 1000S). For GFAP and IBA-1 staining, slices were incubated in H_2_0_2_ (0.3% in PBS) two times for 10 min to remove endogenous peroxidase. For all staining (GFPA, IBA-1, 2E9), slices were then incubated with tween solution (0.3% in PBS) for 10 min to permeabilize tissues and then incubated with blocking solution normal goat serum (2.5% in PBS) for 2h at room temperature. Incubation of primary antibodies diluted in blocking solution was performed overnight at 4°C with the following antibodies 2E9 (specific to human AETA; Sigma-Aldrich, #MABN2295, rat IgG, 2 μg/mL), anti-mouse GFAP (1/400; Sigma-Aldrich, G3893) or anti-rabbit IBA-1 (1/800; Wako 019-19741). Slices were washed 3 times with PBS. For GFAP and IBA-1 staining, slices were then incubated 1h with secondary antibody HRP in PBS 1h at room temperature: anti-mouse (pk-4010, Vector) or anti-rabbit (pk-4001, Vector) and HRP staining was performed with (kit DAB sk-4100, Vector) as described in manufacturer instructions. Slices were washed again 3x in PBS, dried and mounted in Entellan (Sigma-Aldrich, 1.07960). Pictures were taken under a DMD108 Leica microscope at magnification 63x for counting cell number. Quantification of GFAP and IBA-1 positive cells was done using Image J with experimenter blind to genotype during analysis. For 2E9 staining, slices were incubated 1h with secondary antibody anti-rat Alexa-488 (1/1000; Thermofisher; A21208) in PBS 1h at room temperature, washed again 3x in PBS, incubated 10 min with DAPI (1μg/ml; Roche,10236276001), dried and mounted in Mowiol (Sigma-Aldrich, 81381). Pictures where taken under epifluorescence Zeiss Axioplan2.

### qPCR

RNA was extracted from human hippocampi obtained from the NBB (Supplemental Table S1) using the RNeasy Lipid Tissue kit (Qiagen) according to the manufacturer’s instructions. The samples were retrotranscribed using Promega GoScript Reverse Transcriptase kit according to the manufacturer’s instructions in a Biometra TAdvanced thermal cycler according to the program: 5min at 25°C, 50min at 42°C and 15min at 70°C. The qPCR amplification of genes was performed on a mix consisting of 1 μL of RT product diluted at 900ng/µL and 19 μL of mix (containing 10 μL of LightCycler 480 SYBR green I master, 7 μL of UltraPure water, 1μL of forward primer and 1 μL reverse primer). The qPCR was performed with the LightCycler 480 (Roche) (95°C for 5 minutes, (95°C for 10 seconds, 60°C for 30 seconds, 72°C for 30 seconds) x 40 cycles). The melting curve was obtained by increasing the temperature by 1°C every 30 seconds from 70°C to 95°C. Glyceraldehyde-3-phosphate dehydrogenase (GAPDH) was used as a reference gene. Primers used for this qPCR analysis are described in Supplemental Table S3.

Mouse hippocampi were dissected and snap frozen in liquid nitrogen. The samples were retrotranscribed in a Biometra gradient T-type thermal cycler (Applied Biosystems VeritiPro) according to the program: 10sec at 25°C, 120min at 37°C, 5sec at 85°C. The qPCR amplification of genes was performed using SYBR green (Applied Biosystems; #4367659). The samples were diluted to 1/10 (for an RT of 500 ng/20 μL). qPCR was performed on a mix consisting of 2 μL of RT product diluted and 8 μL of mix (containing 5 μL of SYBR green, 2.8 μL of RNase-free water, 0.1 μL of forward primer and 0.1 μL reverse primer). The qPCR was performed with the StepOnePlus Real-Time PCR System (Biosystems) (50°C for 2 minutes, 95°C for 10 minutes, (95°C for 15 seconds, 60°C for 25 seconds) x 40 cycles, 95 ° C for 15 seconds). The melting curve was obtained by increasing the temperature by 1°C every minute from 60°C to 95°C. Peptidylprolyl isomerase A (PPIA, also known as Cyclophilin A) was used as a reference gene. Primers used for this qPCR analysis are described in Supplemental Table S3.

### RNA sequencing

Mouse hippocampi were rapidly dissected and snap frozen in liquid nitrogen. Messenger RNAs (mRNA) from tissues were prepared following the Illumina® Stranded mRNA Prep protocol. They were then sequenced on an Illumina® NextSeq 2000 sequencer, each library receiving at least a 40M reads sequence coverage. Generated FASTQ files were used as input of the nf-core/rna-seq nextflow pipeline (v3.8.1), which performed quality control, trimming and alignment to the mouse reference genome GRCm39 with default parameters (alignment with STAR v2.7.10a followed by quantification with Salmon v1.5.2). Raw counts matrix was then loaded and processed in R (v4.3.1) to perform differential expression analysis following the standard DESeq2 framework (v1.40.2). Genes with a log2FC > 0.2 or log2FC < −0.1, an adjusted P-value < 0.05 and a minimal base mean > 10 in either AETA-m or WT were designated as differentially expressed (DESeq2 analysis output provided in Supplemental Tables S4 and S5 for female data and Table S7 for male data). Functional annotation of genes identified as differentially expressed was performed using the Ingenuity Pathway Analysis (IPA) tool (QIAGEN Inc., https://digitalinsights.qiagen.com/IPA). For comparison with the human NBB-HPC-AD cohort, we extracted the RNA-seq data from the supplemental tables provided by Van Rooij et al. [38]. To compare the neuron function-linked signature obtained in the AETA-m mouse model with the other human AD RNA-seq data, we extracted Supplemental tables provided by Marques-Coelho et al. [26], which correspond to the differential expression analysis between disease and control RNA-seq samples from MSBB BM36 and MSBB BM22 cohorts [48], and the ROSA cohort [9], performed with DESeq2. For comparison with the NBB-PD cohort [7], we ran the differential expression analysis between Braak Lewis Body stages 5 versus 0 (no pathology) samples with the same workflow used for the analysis of our mouse data, starting from the raw count matrix extracted from GSE216281.

### Immunoblotting of p38 and tau in mouse hippocampi

For p38 immunoblotting, freshly dissected mouse hippocampi were homogenized with a potter and then passage through a 1ml syringe with needle (26G) in 150 μL Tris buffer (pH 7.5) containing 2% SDS, 1% triton X100, 10% glycerol, protease inhibitors (#P5726, Sigma-Aldrich) and phosphatase inhibitors (#P8340, Sigma-Aldrich). Lysates were centrifuged for 10 min at 9000 rpm. Proteins in the supernatant were quantified by BCA assay (Pierce) and stored at −80°C until further use. 10 or 20 μg of samples were loaded on precast Novelx 4-20% Tris-tricin gels, in quadruplicates, and transferred to nitrocellulose membranes (0.22 μm, GE Healthcare) which were subsequently incubated with the blocking solution containing 0.2% I-Block (Thermo Fisher Scientific) and 0.1% Tween 20 (Merck) in PBS for 1 hour, followed by overnight incubation with primary antibody in the blocking solution. One set of gels was first immunoblotted for P38, then membranes were stripped, reblocked as above and then immunoblotted for P-P38. Another set of gels was first immunoblotted for P-P38 and then P38. The stripping procedure consisted of incubating membranes in ddH_2_0 containing 1.5g/L glycine, 0.1% SDS, 1% tween 20 for 15 min at room temperature followed by rinsing two times for 5 min with PBS. Primary antibodies used were rabbit anti-MAPK p38 (#8690, Ozyme; diluted 1/1000) and mouse anti-phospho-p38 (#9216, Ozyme; diluted 1/400). Antibody detection was performed using the corresponding anti-mouse/rabbit-IgG-HRP conjugated secondary antibody (Thermo Fisher Scientific) and chemiluminescence detection reagent ECL (Thermo Fisher Scientific). Quantification of immunoblots to calculate P-p38/p38 ratio was performed using ImageJ software.

For Tau and phosphorylated tau immunoblotting, previously snap-frozen mouse hippocampi were homogenized in 200 μl Tris buffer (pH 7.4) containing 10% sucrose and protease inhibitors (Complete; Roche Diagnostics) and sonicated. Homogenates were kept at −80°C until use. Protein concentrations of the samples were quantified using the BCA assay (Pierce), diluted in lithium dodecyl sulphate buffer supplemented with reducing agents (Invitrogen) and then separated on 4-12% Criterion Bis-Tris Gels (Invitrogen). Proteins were transferred to nitrocellulose membranes, which were then saturated with 5% non-fat dried milk in TNT (Tris 15mM pH 8, NaCl 140mM, 0.05% Tween) and incubated at 4°C for 24h or 48h with the primary antibodies. Rabbit polyclonal anti-C ter (home-made, D. Blum, 24h incubation) was diluted at 1/4000. Mouse monoclonal anti-Tau1 (Millipore # #MAB3420, 24h incubation) was diluted at 1/10000. Rabbit polyclonal anti-phospho-Tau (Ser199) (home-made, 24h incubation) was diluted at 1/2000. Rabbit polyclonal anti-phospho-tau (Ser396) (Invitrogen #44-752G, 48h incubation) was diluted at 1/10000. Appropriate HRP-conjugated secondary antibodies were incubated for 45 min at RT. Signals were visualized using chemiluminescence kits ECL (#RPN2106; Amersham) or ECL Prime (#RPN2232; Amersham) and an Amersham ImageQuant 800 imaging system. Quantifications were performed using ImageJ software and results were normalized to β-actin for C-ter and Tau-1 levels, and normalized to β-actin and to C-ter Tau levels for phosphorylated Tau pS199 and pS396.

### Electrophysiology in hippocampal slices

For ex vivo electrophysiology recordings, 6-10 months old males and females AETA-m and WT littermates were used. Mice were culled by cervical dislocation. Hippocampi were dissected and sliced (250 μm for patch-clamp recordings, 350 μm for field recordings) on a vibratome (Microm HM600V, Thermo Scientific, France).

Different external solutions for slicing, recovery and recording were used in this study as follows. For most LTP recordings (except experiment in Figure S4c-d), after dissection, hippocampi were incubated for 5 min and then sliced in ice-cold oxygenated (95 % O_2_/ 5 % CO_2_) cutting solution (in mM): 234 sucrose, 2.5 KCl, 1.25 NaH2PO4, 10 MgSO4, 0.5 CaCl2, 26 NaHCO3, 11 glucose (pH 7.4). For recovery (1h at 37°C) and recordings (at 27-29°C), slices were then incubated in artificial cerebral spinal fluid (aCSF; in mM): 119 NaCl, 2.5 KCl, 1.25 NaH2PO4, 26 NaHCO3, 1.3 MgSO4, 2.5 CaCl2 and 11 D-glucose, oxygenated with 95 % O2 and 5 % CO2, pH 7.4 for 1h at 37 ± 1°C. For field LTD recordings, after dissection, hippocampi were incubated for 5 min, sliced in ice-cold oxygenated (95 % O_2_/ 5 % CO_2_) and then incubated for recovery (1h at 37°C) in the same condition as described above for LTP. To optimize LTD recordings, slices were recorded (at 27-29°C) in another aCSF (in mM): 119 NaCl, 2.5 KCl, 1.25 NaH2PO4, 26 NaHCO3, 2 MgSO4, 4 CaCl2 and 11 D-glucose, oxygenated with 95 % O2 and 5 % CO2, pH 7.4 for 1 h at 37 ± 1 °C, with picrotoxin (50 μM; Sigma-Aldrich, France) supplementation to block GABA_A_ receptors. For experiment in Figure S4c-d, after dissection, hippocampi were incubated for 5 min and then sliced in ice-cold oxygenated (95 % O_2_/ 5 % CO_2_) cutting solution (in mM): 206 sucrose, 2.8 KCl, 1.25 NaH2PO4, 2 MgSO4, 1 CaCl2, 26 NaHCO3, 0.4 sodium ascorbate, 10 glucose (pH 7.4). For recovery (1h at 37°C) and recordings (at 27-29°C), slices were then incubated in aCSF (in mM): 124 NaCl, 2.8 KCl, 1.25 NaH2PO4, 26 NaHCO3, 2 MgSO4, 3,6 CaCl2, 0.4 sodium ascorbate and 10 D-glucose, oxygenated with 95 % O_2_ and 5 % CO_2_, pH 7.4 for 1h at 37±1°C.

For all whole-cell patch-clamp recordings, slicing, recovery and recordings were made in same aCSF as for most LTP experiments described above. Recordings were performed at 30-31°C. The recording aCSF was supplemented with picrotoxin (50 μM; Sigma-Aldrich, France). For NMDAR spontaneous excitatory post-synaptic current (sEPSC) recordings, the aCSF was also supplemented with NBQX (10 μM; Tocris, England) to block AMPAR activity. For AMPAR sEPSC recordings, the aCSF was also supplemented with APV (50 μM; Tocris, England) to block NMDAR activity.

All recordings were made in a recording chamber on an upright microscope with IR-DIC illumination (SliceScope, Scientifica Ltd, UK) using a Multiclamp 700B amplifier (Molecular Devices, San Jose, CA, USA), under the control of pClamp10 software (Molecular Devices, San Jose, CA, USA). Data analysis was executed using Clampfit 10 software. Field excitatory post-synaptic potentials (fEPSPs) were recorded in the stratum radiatum of the CA1 region (using a glass electrode filled with 1 M NaCl and 10 mM 4-(2-hydroxyethyl)-1-piperazineethanesulfonic acid (HEPES), pH 7.4) and the stimuli were delivered at 0.1 Hz to the Schaffer collateral pathway by a monopolar glass electrode filled with aCSF. fEPSP response was set to approximately 30% of the maximal fEPSP response i.e. approx. 0.2–0.3 mV, with stimulation intensity 10 μA ± 5 μA delivered via a stimulation box (ISO-Flex, A.M.P.I. Inc., Israel). A stable baseline of 20 min was first obtained before induction for long-term plasticity recordings. The time courses were obtained by normalizing each experiment to the average value of all points constituting a 20 min stable baseline before induction. fEPSP magnitude was measured during the last 15 min of recording (45–60 min after induction) and calculated as % change fEPSP slope from baseline average.

For patch clamp recordings, after obtaining a tight seal (>1GΩ) on the cell body of the selected neuron, whole-cell patch clamp configuration was established, and cells were left to stabilize for 2-3 min before recordings began. Holding current and series resistance were continuously monitored throughout the experiment, and if either of these two parameters varied by more than 20%, the cell was discarded.

For paired-pulse ratios (PPR), EPSCs were obtained and two stimuli were delivered at 100, 200, 300 and 400 ms inter-stimulus interval (ISI). PPR was calculated as EPSC2 slope/EPSC1 slope (10 sweeps average per ISI). Recordings of WT and AETA-m mice were interleaved between days. Internal solution was a CS-gluconate solution: 117.5 mM Cs-gluconate, 15.5 mM CsCl, 10 mM TEACl, 8 mM NaCl, 10 HEPES, 0.25 mM EGTA, 4 mM MgATP and 0.3 NaGTP (pH 7.3; osmolarity 290-300 mOsm).

AMPAR sEPSCs were recorded at −65 mV using the same CS-gluconate internal solution detailed above. NMDAR sEPSCs were recorded at −65mV using a Cesium-methanesulfonate internal solution (mM): 143 Cesium-methanesulfonate, 5 NaCl, 1 MgCl2, 1 EGTA, 0.3 CaCl2, 10 Hepes, 2 Na2ATP, 0.3 NaGTP and 0.2 cAMP (pH 7.3 and 290-295 mOsm). sEPSCs were recorded in gap-free mode for 5 min for individual neurons of each genotype (WT and AETA-m). The Clampfit 10.6 software (Axon instruments) was used for analysis of sEPSCs to compare frequencies and amplitudes. sEPSCs were detected manually by following criteria of peaks with a threshold 2xSD of baseline noise level and a faster rise time than decay time. Analyses were performed blind to experimental condition. In the NMDAR sEPSC condition, we had previously ensured that the recorded currents were NMDAR current as they were absent in presence of APV [11].

### Golgi Cox staining for spine density analysis

Brains were rapidly collected and impregnated in a Golgi-Cox solution (1% potassium dichromate, 1% mercuric chloride, 0.8% potassium chromate) for 3 weeks at room temperature according to manufacturer instruction (FD rapid GolgiStain kit, FD Neurotechnologies, USA). Brains were sectioned coronally (80 µm) using a vibratome, and stained and mounted according to protocol. Images were acquired under white light on a DMD108 Leica microscope (60X magnification). Spine density (number of spines per 1 µm length) in selected segments (minimum of 30 per mouse) of secondary dendrites of CA1 pyramidal neurons in stratum radiatum was estimated using Image J and manual count by an experimenter blinded to mouse genotype. As several batches of female mice were processed at independent times (always pairing WT with AETA-m mice), the values per batch were normalized to the average WT spine density value before pooling the data for final statistical analysis per mouse. As only one batch of male mice was processed, we did not require this additional normalization and data are thus presented as spine number/μm.

### Morris Water Maze

AETA-m (6-10 months old) were submitted to the Morris water maze task in either a 90 cm or a 150 cm pool. The apparatus consisted in a circular tank (Ø 90□cm or 150 cm) filled with water (temperature 25□±□1□°C) made opaque with the addition of white opacifier (Viewpoint, France). For the 90 cm pool protocol, the test consisted in 3 phases: (1) Cue task training (2□days), (2) Spatial learning training (7□days) and (3) Long-term reference memory Probe test (at 24h and at 7 days). An escape platform (Ø 8□cm) was submerged 1□cm below the water surface for the cue task training and the spatial learning training. The animals performed 4 trials/day with a maximum trial duration of 90s (+30s on the platform at the end of each trial) and with an inter-trial interval of 10 min. During the cue task, a visible flag was placed on the top of the submerged platform and the tank was surrounded by opaque curtains. The mice were allowed to find the visible platform. The platform emplacement was changed for the second day of the cue task (Day 2). For the spatial learning training phase, extra-maze cues were placed on the walls surrounding the maze to allow the animals to create a spatial map and find the submerged platform. A new platform placement was designed and kept in the same position for the 7 training days. Probe test was run to assess the strength of the memory for the platform location: the platform was removed, and the test mouse was allowed to search for it for 90s. For the 150 cm pool protocol, the test consisted in 3 phases: (1) Cue Task training (1 day), (2) Spatial Learning training (8 days) and (3) long reference memory Probe test (at 24h and at 7 days (for males only)). Similarly, an escape platform (Ø 8□cm) was submerged 1□cm below the water surface for the cue task training and the spatial learning training. During Cue Task training, a visible flag was placed on top of the submerged platform and the tank was surrounded by opaque curtains. Mice were allowed to find the visible platform. The mice performed the test along 2 sessions (T1 and T2) of 3 trials each with a maximum trial duration of 90s (+30s on the platform at the end of each trial) and with an inter-trial interval of 10 min. For the Spatial Learning training phase, extra-maze cues were placed on the walls surrounding the maze to allow the animals to create a spatial map to find the submerged platform and the tank was surrounded by white and patterned curtains. A new platform placement was designed and kept in the same position for 8 training days until the WT group reached the performance criteria. The probe test was run at 24h and 7 days (for males only) to assess memory strength at which time the platform was removed, and the test mouse was allowed to search for it for 60s. For both protocols, the training phase was stopped once the WT mice exhibited an average latency that was statistically different from the first day of training and not statistically different from 20s. All the trials were video recorded and tracked using ANYmaze software (Stoelting, Wood Dale, USA) to analyze escape latency.

### Statistical analysis

Results are shown as mean ± s.e.m. Numbers and their correspondence are given in each figure. Statistical analysis was performed with GraphPad Prism 9 software (Dotmatics). Statistical analyses are described in brief in figure legends and are presented in detail in Supplemental Tables S8-S16 (one table per figure). Statistical significance was set at p < 0.05; *p < 0.05; **p < 0.01; ***p < 0.001; ****p < 0.0001.

### Use of AI and AI-assisted technologies

ChatGPT (OpenAI, USA) was used in order to improve readability and language. After using this tool, the lead author reviewed and edited the content as needed and takes full responsibility for the content of the publication.

### Availability of data and materials

Raw and processed RNA-seq data will be deposited on in the Gene Expression Omnibus.

## Results

### AETA levels are increased in AD patient brain

AETA levels were quantified in hippocampi and prefrontal cortices (PFC) of AD patient brains (Braak stage V/VI) against age-matched control subjects (Supplemental Table S1 and S2) using protocols and antibodies largely characterized in mouse and human samples as described previously [51]. We extracted the soluble protein fraction from these brains (DEA fraction). Control and AD brain DEA fractions displayed preserved protein integrity as exemplified by GAPDH protein content (Supplemental Figure S1a). We observed a significant increase in AETA in both hippocampi and PFC of AD brains as compared to controls (Figure 1b; Supplemental Figure S1b). As expected, AETA was represented as a multiband signal [51]. This increase in AETA levels was not due to changes in protein (Figure 1c) or mRNA expression (Supplemental Figure S1c) of APP. We also calculated the AETA/APP ratio and observed an increase of this ratio (Supplementary Figure S1d), confirming that the increase in AETA levels in hippocampi of human AD brains is not due to an increase in APP levels *per se*. We also compared AETA levels between both genders in hippocampi in either control subjects or AD patients. While AETA levels were similar in hippocampi of male and female control subjects (Supplementary Figure S1e), they were increased in AD female when compared to AD males (Figure 1d). Together, these data show that AETA is increased in brains of human AD patients, with a stronger increase in women.

### Chronic increase in AETA brain levels leads to human AD-like astrocyte and microglia reactivity and a molecular signature of synapse dysfunction in females

Considering this increase in AETA levels in brains of AD patients, we asked if a chronic increase in brain AETA levels could contribute to AD pathology. To address this question, we used a new mouse model, named AETA-m, which harbors a transgene allowing expression of a secreted form of human AETA (long α form) under the Thy1.2 neuronal promoter (Figure 2a). In this model, human AETA expression was observed at similar levels between 1 and 12 months of age, albeit with some inter-individual differences. Notably, no accumulation of human AETA was evident with age in this model (Figure 2b). We quantified the average amount of human AETA being expressed in hippocampi of these mice by comparing to known amount of synthetic human AETA peptide. Human AETA concentration was calculated as approximately 0.27 ng/μg of protein (Supplemental Figure S2a). We had confirmed secretion of the peptide, prior to generation of the mouse model, by transfecting the original cDNA (pSecTag 2A-human AETA with a myc-flag tag) in CHO cells and immunoblotting the medium of these cells (Supplementary Figure S2b). Further, we also confirmed secretion of this peptide by detecting its presence in the cerebrospinal fluid (CSF) extracted from the cisterna magna of female AETA-m mice (Supplementary Figure S2b). We did not detect any difference in levels of the peptide when quantified in hippocampi of adult male and female mice (Supplementary Figure S2c). We also analyzed this new mouse model for other parameters to check for general health. Body weights at weaning and in adult AETA-m mice were normal (Supplemental Figure S2d). Human AETA expression was strong in the CA1 and CA3 regions of the hippocampus and to a lesser extent in the dentate gyrus (Supplementary Figure S2e). The general cytoarchitecture of the hippocampus was normal as measured by the thickness of the CA1 cell body layer in hippocampi slices stained with cresyl violet (Supplemental Figure S2f), suggesting normal neuron density. This expression did not alter levels of endogenous mouse AETA or endogenous Aβ_1-40_, as published in a preliminary analysis of hippocampal protein content from these AETA-m mice [11]. We further analyzed the levels of endogenous full-length APP and other peptides associated to APP processing (CTFη and sAPPα/β). Levels of these endogenous peptides were unchanged (Supplemental Figure S2g) showing that increasing AETA levels does not alter endogenous APP processing in these mice. We also tested the neuroinflammatory status of brains of AETA-m mice by analysis the number of astrocytes and microglia cells, by counting the number of GFAP+ and IBA-1+ positive cells respectively, in the CA1 region of the hippocampus. We observed an increase in the number of both cell types in females, but not in male mice at 7 months of age (Figure 2c-d), suggesting that an increase of AETA in female mice led to proliferation of these cell types. We next asked if this proliferation was correlated to an increase in neuroinflammatory markers. We tested the mRNA levels of a set of genes known to be increased when neuroinflammation is fully developed. We quantified mRNA levels of Trem2, C1qa, ccl4 and Vim genes. (Supplemental Table S3). We could not detect any alterations in the transcription of any of these genes in either sexes (Figure 2e), suggesting that AETA-m mice do not display a strong neuroinflammatory status. We also tested the status of Tau phosphorylation in hippocampi of these mice to seek for this additional hallmark of AD pathology [1, 18, 27]. We could not detect any evidence of tau hyperphosphorylation (Supplemental Figure S3).

**Fig. 2.**
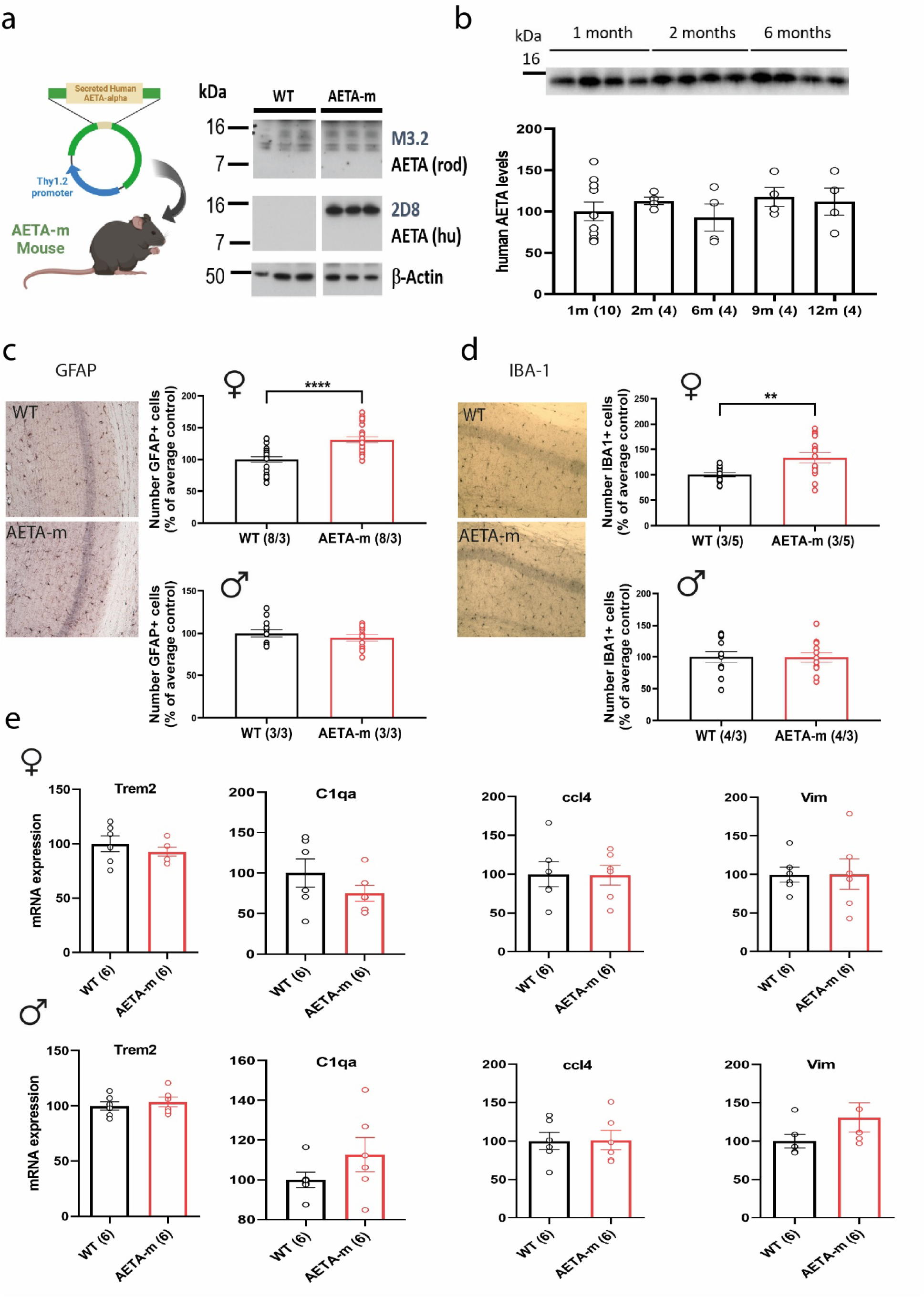
Neuroinflammatory status of new AETA-m mouse model. **(a)** Diagram of new AETA-m mouse model expressing secreted human AETA with example of immunoblot to detect endogenous AETA (rodent (rod); M3.2 antibody) and recombinant secreted human (hu) AETA (long AETA-alpha form) (2D8 antibody) and β-actin (3 wild type (WT) and 3 AETA-m mice). **(b)** Example of immunoblot with 2E9 antibody and quantification of recombinant human AETA in hippocampi of male AETA-m mice at different ages (1-12 months; n= mice). Full blots are provided in Supplemental material. **(c)** Number of GFAP+ cells in CA1 region of hippocampus in 7 months old female and male mice (normalized to average control) (n=slice/mouse number analyzed). **(d)** Number of IBA-1+ cells in CA1 region of hippocampus in 7 months old female and male mice (normalized to average control) (n=slice/mouse number analyzed). **(e)** mRNA expression levels, quantified by qPCR, of markers of neuroinflammation (Trem2, C1qa, ccl4 and Vim) in 6-8 months old female and male WT and AETA-m mice. n= mice for b and e; N/n= slices/mice for c-d. Error bars represent s.e.m. ** p < 0.01; **** p < 0.0001. Statistics: one-way ANOVA for b, Student t-test for c-e (except for Mann-Whitney for male Vim analysis). See full statistics in Supplemental Table S10.

To broadly address the molecular changes promoted by the production of AETA in the hippocampus, we performed RNA sequencing of hippocampi from female adult AETA-m mice and WT littermates. We chose female mice for this analysis as this sex displayed higher AETA levels in human AD brains and exhibited astrocyte and microglia reactivity. We identified 567 differentially expressed genes between both conditions (Supplemental Table S4 and S5). Within this dataset, we focused our attention to a major hallmark of AD that is neuronal dysfunction, notably of synaptic pathways linked to NMDA receptor activity [22, 24, 40]. Functional annotation of this molecular signature with IPA revealed modifications in diseases & molecular functions, signaling pathways, and upstream regulators tightly linked to neuron function (Figure 3a, Supplemental Table S6), with all these pathways showing mild but significant downregulated expression in AETA-m mice, except for ‘ion channel transport’ which was selectively upregulated. Notably, most genes associated to ‘SNARE signaling pathway’ and ‘Assembly and cell surface presentation of NMDA receptors’ with IPA, which are particularly relevant to synapse function, were significantly down-regulated in hippocampi of AETA-m mice (Figure 3b). To identify possible sexual dimorphisms in this transcriptional adaptation in the AETA-m line, as evidenced for astrocyte and microglia reactivity, we also performed bulk RNAseq for hippocampi of male AETA-m and WT littermates (Supplementary Table S7) and searched for this synaptic molecular signature. Interestingly, there were no modifications of transcription of these genes (Figure S4a), suggesting that male and female hippocampi adapt differently at the transcriptional level in the presence of chronically increased AETA. Of note, making use of these RNAseq data, we also analyzed an additional panel of genes pertinent to neuroinflammation and did not observe any significant alterations in either sex (Supplementary Figure S4c). confirming our qPCR data (Figure 2e).

**Fig. 3.**
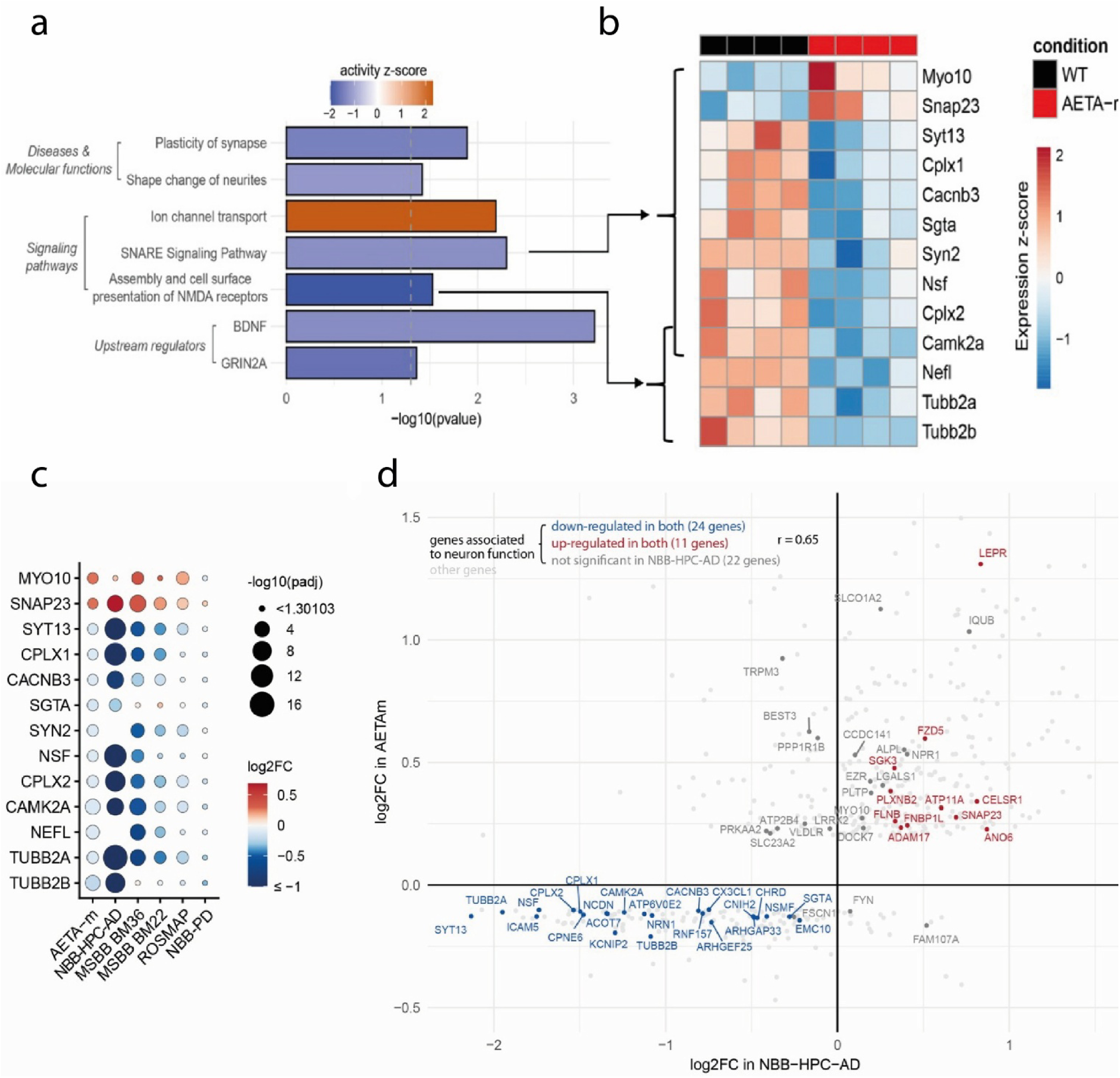
Modifications of transcriptome of signaling pathways relevant for neuron function of hippocampus of female AETA-m mice resemble human AD signature. **(a)** Results of Ingenuity Pathway Analysis focusing on differentially expressed neuron-relevant pathways identified by RNA-seq data analysis of 4 AETA-m and 4 WT 6-8 months old female mice. The IPA z-score indicates whether the activity of each pathway is predicted to be increased (z-score > 0) or decreased (z-score < 0) in AETA-m mice. Genes belonging to these pathways are listed in Supplemental Table S6. **(b)** Heat map of normalized and scaled expression of genes associated to “SNARE signaling pathway” and “Assembly and cell surface presentation of NMDA receptors” in the 8 sequenced samples. **(c)** Comparison of differential expression (relative to respective controls) of genes highlighted in (b) from hippocampi of AETA-m mice to human NBB-HPC-AD, MSBB BM36, MSBB BM22, ROSMAP and NBB-PD RNA-seq datasets. **(d)** Correlation between the log2FC of the 540 differentially expressed genes identified in hippocampi of the AETA-m model (Supplemental Tables S4 and S5) and their log2FC obtained from the analysis of NBB-HPC-AD RNA-seq dataset. Highlighted are the 66 genes associated to neuron function using IPA (as shown in (b) and listed in Supplemental Table S6).

Next, we sought to determine if this molecular synaptic signature observed in AETA-m female mice was pertinent to human AD pathology, particularly the hippocampus and para-hippocampal regions, which are most vulnerable in AD. We compared the synaptic transcriptional signature obtained in female AETA-m hippocampi (Figure 3c) with publicly available transcriptomic data of Van Rooij et al. [38] where they compared bulk RNAseq data of hippocampi of 20 AD cases and 10 age- and sex- matched cognitively healthy controls (NBB-HPC-AD). We observed a strong correlation with most mRNA being up and down-regulated in both set of data (Figure 3c). Indeed, 11 out of the above-mentioned 13 genes associated to the to ‘SNARE signaling pathway’ and ‘Assembly and cell surface presentation of NMDA receptors’ pathways also showed significant expression differences compared with control brains and displayed identical up (two genes) or down (nine genes) regulation (Figure 3c), the other two genes (SYN2 and NEFL) not being reported in the human RNAseq data analysis. We also compared the AETA-m female data to the female NBB-HPC cases only. Although there were fewer human cases to compare with (5 Controls vs 6 AD female cases) thus reducing the statistical power of the alterations observed in this cohort, we could still observe this signature (Supplementary Figure S4b). Using publicly available transcriptomic data of the Mount Sinai AD cohort (MSBB) [48], we also compared the synaptic signature obtained in female AETA-m mice with data bulk RNAseq data obtained from the human Brodmann area 36 (BM36). This parahippocampal area is highly relevant to formation, consolidation and retrieval of hippocampus-linked memories [8] and particularly vulnerable, even amongst temporal lobe regions, in human AD brains showing the most profound molecular alterations as previously evidenced with microarray profiling [49]. A molecular signature similar to that observed in AETA-m hippocampi was also evident in this region. We also compared the AETA-m molecular signature to another temporal lobe area of the MSBB cohort (Brodmann area 22 (BM22)), which was defined as a region less vulnerable than BM26 in AD [49]. This molecular signature was slightly less pronounced, although 9 out of 13 genes exhibited a similar pattern (Figure 3c). We also tested if this molecular signature could be evidenced in another publicly available dataset from a different cohort, The Religious Order Study and Memory and Aging Project (ROSMAP) dataset, where the transcriptome profiling was performed on the dorsolateral prefrontal cortex of control and AD post-mortem tissue [9]. This brain region is highly implicated in the control of executive functions, including working memory and cognitive flexibility, both of which are impaired during AD progression [20]. The molecular signature was less obvious in this dataset, but still 7 out of the 13 genes displayed a similar pattern to the molecular signature of AETA-m hippocampi (Figure 3c). To confirm a specific link of this molecular signature to AD, we also compared our mouse data to a publicly available dataset from frontal cortex samples of Parkinsonian human patients (NBB-PD) [7]. In this dataset, we used the differential profiling observed between Braak Lewis Body stages 0 (no pathology) and 5 (moderate Lewis Body pathology in frontal cortex). This molecular signature was completely absent in this PD dataset (Figure 3c). As the NBB-HPC-AD dataset exhibited the strongest similarity to the female AETA-m synaptic signature, we extended the comparison to all genes identified as significantly differentially expressed in the AETA-m model (567 genes on which 540 have a known human ortholog, Supplemental Tables S4 and S5). By correlating the log2FC obtained in both analyses, we obtained a Pearson’s correlation coefficient of 0.52. By selecting only the genes contributing to the neuron-relevant enriched pathways highlighted in Figure 3a and in Supplemental Table S6, this correlation reached a coefficient of 0.65 (Figure 3d). It is worth noting that the synaptic genes (in blue in Figure 3d) exhibited a stronger down-regulation in the human NBB-HPC cohort than in female AETA-m mice. This could be due to a more advanced synapse deterioration in human AD brains, all at Braak stages V-VI, while AETA-m mice probably display early synapse alterations without overt neuronal loss. Together, these data demonstrate that the neuronal molecular signature, and particularly the synaptic signature, observed in hippocampi of female AETA-m mice is reminiscent of human AD pathology with the strongest correlation observed in memory-encoding regions, which are highly vulnerable in AD.

### Increased brain AETA levels modify NMDA receptor-linked signaling

We next tested if, in AETA-m mice, these molecular alterations correlated with hippocampal synapse dysfunction by analyzing NMDAR-dependent long-term potentiation (LTP) and long-term depression (LTD) at the CA3-CA1 synapse in acutely dissected hippocampal slices of adult (5-10 months) AETA-m and WT littermates. We observed that LTP was reduced in both female and male AETA-m mice (Figure 4a and Supplemental Figure S5a). We observed enhanced LTD at this same synapse in male AETA-m mice (Figure 4b). LTD in WT female mice could not be induced at this age using this NMDAR-dependent LTD induction protocol (Supplemental Figure S5b). Plasticity deficits were postsynaptic as presynaptic plasticity, measured by the paired-pulse ratio, was normal at this synapse (Supplemental Figure S5c). We tested both AMPA receptor (AMPAR) and NMDAR signaling, the two receptors required for these plasticity mechanisms [23], by recording spontaneous excitatory synaptic currents (sEPSC) in CA1 pyramidal neurons. AMPAR signaling, as measured by AMPAR sEPSC frequency and amplitude, was not affected (Figure 4c), while NMDAR sEPSC frequency, but not amplitude, was reduced (Figure 4d). Lower LTP and enhanced LTD could be explained by compromised NMDAR signaling, possibly due to inhibition of calcium entry via NMDARs in presence of AETA [11]. To test this possibility, we tested LTP again at this synapse using a different extracellular medium containing higher calcium [51]. We observed that, in this condition, LTP was normal (Supplemental Figure S5d). This suggests that, although NMDAR-dependent signaling is compromised, it might be due to over-activation of AETA-linked signaling that shunts calcium entry via partial inhibition of NMDAR ionotropic activity [11]. To test this further, we examined p38 MAP kinase activation in hippocampi of these mice. This activation was shown to be important for LTD and spine shrinkage via the unconventional non-ionotropic function of NMDARs and favored in presence of AETA [11, 31, 33]. P38 phosphorylation was enhanced in AETA-m hippocampi of both female (Figure 4e) and male mice (Supplementary Figure S5e). This was linked to a reduction of spine density in dendrites of CA1 pyramidal neurons in both female (Figure 4f) and male AETA-m mice (Supplemental Figure S5f). Together these data suggest that, upon chronic increase of AETA levels, NMDAR signaling is particularly compromised favoring synapse weakening both functionally and structurally.

**Fig. 4.**
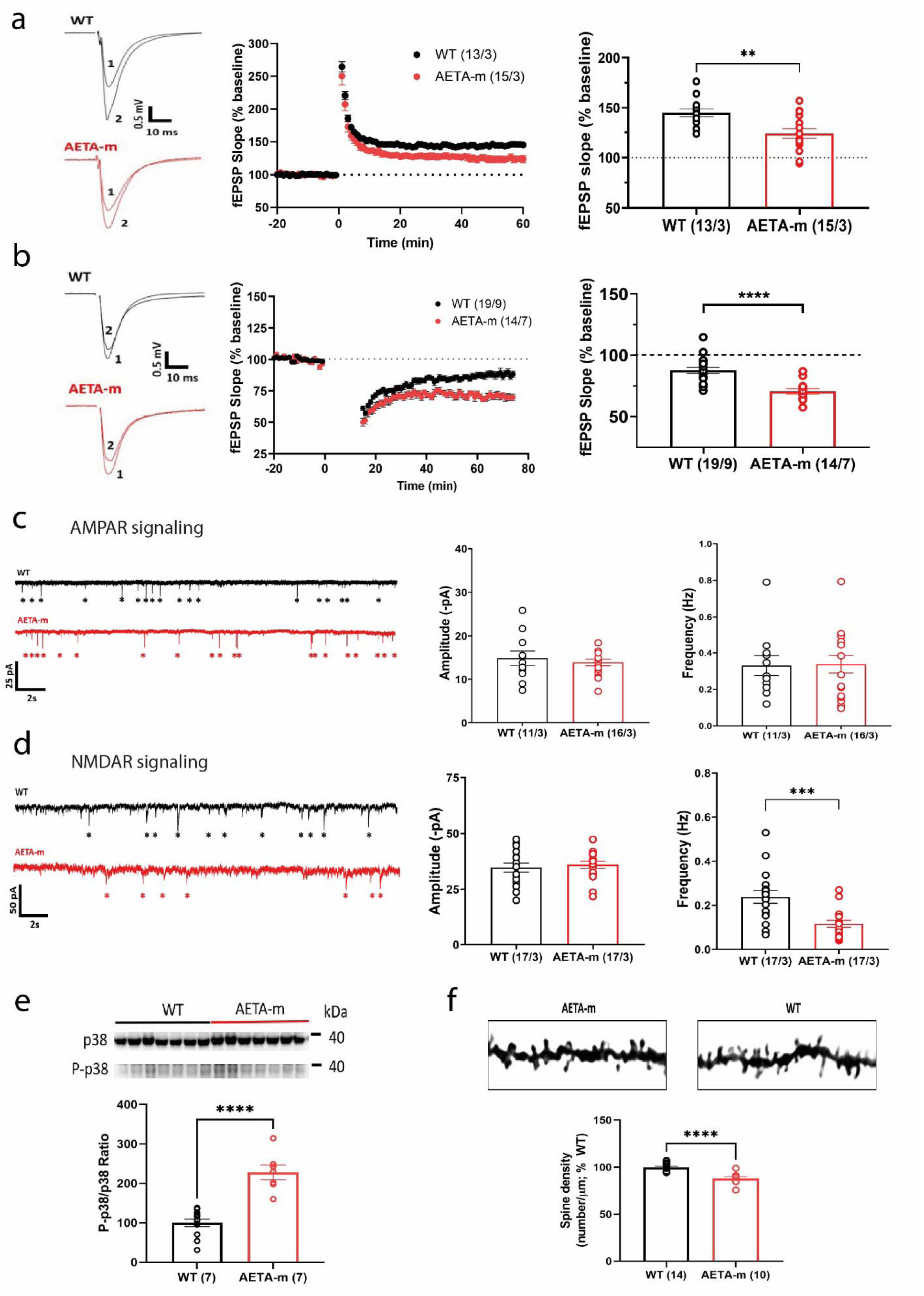
Increased AETA brain levels modify hippocampal synaptic signaling in AETA-m mice. **(a)** Representative traces (left, 1 shows trace pre-, 2 shows trace post-induction), time course (middle; induction at time 0) and bar graph (right, 45-60 min post-induction) of long-term potentiation of fEPSP slope (% baseline) at CA3-CA1 synapse in hippocampal slices of female WT and AETA-m mice. **(b)** Representative traces (left, 1 shows trace pre-, 2 shows trace post-induction), time course (middle; induction at time 0) and bar graph (right, 45-60 min post-induction) of long-term depression of fEPSP slope (% baseline) at CA3-CA1 synapse in hippocampal slices of male WT and AETA-m mice. **(c)** Representative traces (left), amplitude (middle) and frequency (right) of AMPAR sEPSCs in CA1 pyramidal neurons of female WT and AETA-m mice. **(d)** Representative traces (left), amplitude (middle) and frequency (right) of NMDAR sEPSCs in CA1 pyramidal neurons of female WT and AETA-m mice. **(e)** Example immunoblots of p38 and phosphorylated p38 (P-p38) and P-p38/p38 ratio quantification in female WT and AETA-m mice. Samples were tested in quadruplicate and quantified per mouse. **(f)** Example images of segments of dendrites present in the stratum radiatum of CA1 pyramidal neurons from female WT and AETA-m mice stained by Golgi-Cox and quantitative analysis of spine density on these segments. n/N= slices/mice for a-d; n= mice for e-f. Error bars represent s.e.m.; ** p < 0.01; *** p < 0.001; **** p < 0.0001. Statistics: Student t-test for all except for Mann-Whitney test for 4c right. See full statistics in Supplemental Table S13.

### AETA-m mice exhibit memory decay reminiscent of prodromal AD

As hippocampi of AETA-m mice display an AD-like molecular signature of synapse dysfunction in females and NMDAR associated synapse weakening, we would expect hippocampus-dependent memory alterations in AETA-m mice, at least in females. We thus submitted both male and female mice to the Morris water maze task. Mice were first tested in a small Morris water maze pool (90 cm diameter). Two days of cue task was performed to ensure that AETA-m mice exhibited normal locomotor activity and could see visual cues adequately (Figure 5a, Supplemental Figure S6a). Mice were trained until WT mice reached the criterion of less than 20s delay to reach the platform. AETA-m mice exhibited normal learning (Figure 5a, Supplemental Figure S6a). After a 24h delay, memory retrieval was measured as time spent in target quadrant. Memory retrieval was normal in AETA-m mice (Figure 5b, Supplemental Figure S6b). To test for persistence of memory consolidation, we submitted these mice to a second probe test 7 days later. No significant difference was observed between the two genotypes, although, in AETA-m females, the 7-day memory was slightly weaker than WT females when compared to chance level (25%) (Figure 5c and Supplemental Figure S6c). We also analyzed the accuracy of this memory measured as time spent in platform zone and number of platform crossings. While WT mice conserved a stable ability to locate this position between 24h and 7 days, AETA-m mice, most notably female AETA-m mice, exhibited a decay of this accuracy (Figure 5d-e and Supplemental Figure S6d-e). We also challenged AETA-m mice in a larger pool (150 cm), more demanding in terms of memory accuracy, asking if we could exacerbate this memory phenotype. Again, during this new protocol, AETA-m mice performed the cue task correctly and training of AETA-m mice was normal (Figure 5f and Supplemental Figure S6f). After a 24h delay WT and AETA-m females correctly located the target quadrant (their scores were significantly above chance level), but performance of AETA-m females was significantly lower compared to WT females in this larger pool (Figure 5g). Male mice performed as well as WT mice, even for the 7 days probe test (Supplemental Figure S6g-j). These results indicate that female AETA-m mice exhibit weaker consolidation of long term memory, suggesting prodromal human AD-like cognitive deficits [30, 47].

**Fig. 5.**
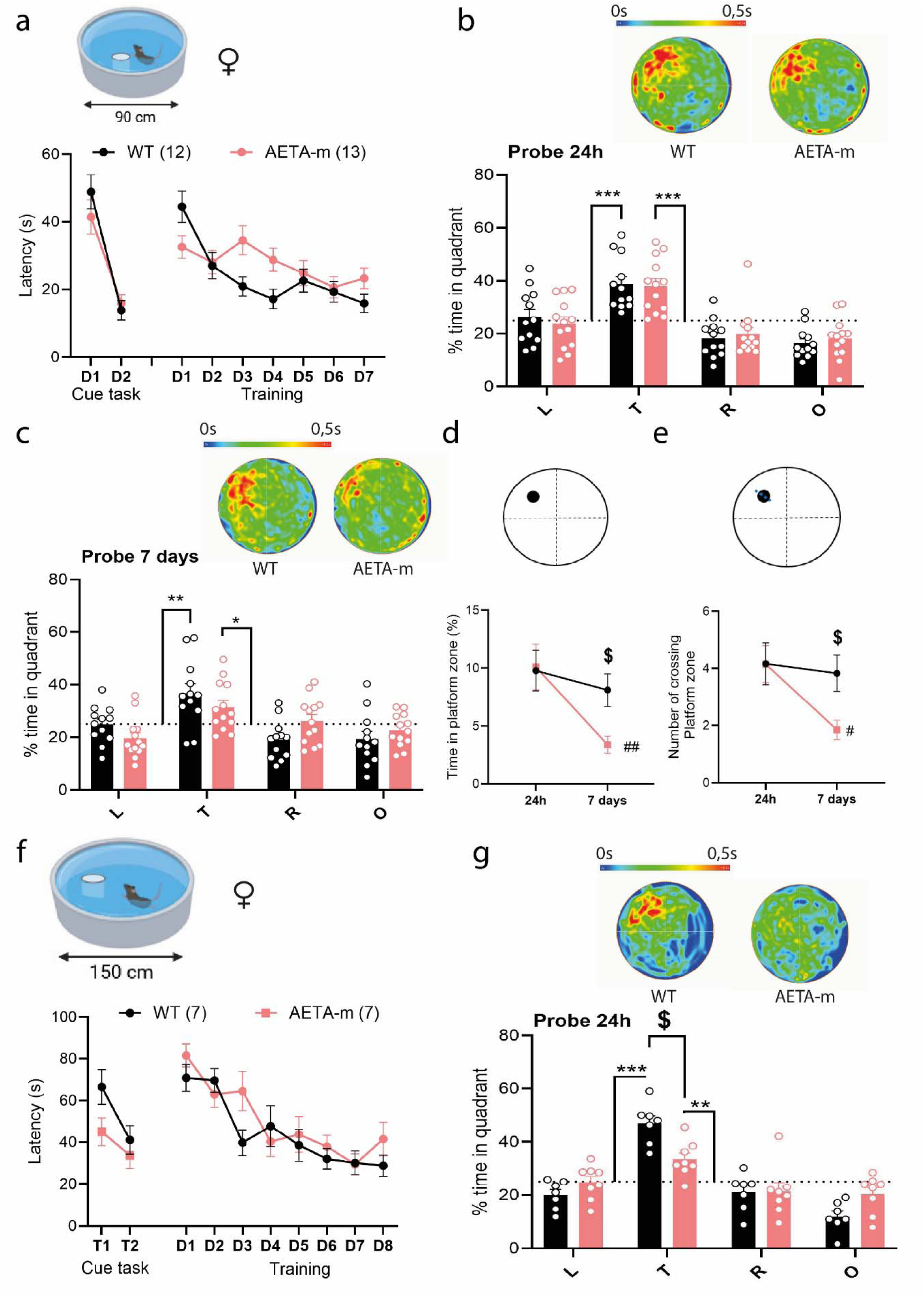
Long-term memory deficits in female AETA-m mice. **(a)** Latency (s) during cue task and training in 90 cm Morris water maze pool. **(b-c)** Average heat maps (top) and % time spend in quadrants (T: target, L: left, R: right, O: opposite) during probe test (b) at 24h, (c) at 7 days. **(d)** Time spent in platform zone (black dot in diagram) at 24h and 7 days. **(e)** Number of platform crossing (dotted line in diagram) at 24h and 7 days. **(f)** Latency (s) during cue task and training in 150 cm Morris water maze pool. **(g)** Average heat maps (top) and % time spent in quadrants during probe test at 24h. n= mice. Error bars represent s.e.m.; *p < 0.01; ** p < 0.001; *** p < 0.0001 against chance level (25%, dotted line on graph); $ p < 0.01 between WT and AETA-m mice; # p < 0.01; ## p < 0.001 between 24h and 7 days within genotype. Statistics: Two-way ANOVA. See full statistics in Supplemental Table S15.

## Discussion

Aβ has been long suspected to be the main culprit of synapse dysfunction in AD. There is strong evidence supporting this claim, although the exact mechanism by which it impairs synapse signaling remains unclear. Yet, an increasing number of studies suggest that other amyloid-independent mechanisms could also lead to synapse dysfunction in AD, beyond Aβ actions [4, 10, 19, 35, 44]. Our study provides the first evidence that AETA could represent an important additional factor of this AD hallmark. Here, we show that AETA levels are elevated in the hippocampus and PFC of AD patients, with sex-dimorphism characterized by a stronger increase in AD female compared to AD male. Increasing AETA levels in the mouse brain leads to hippocampal transcriptomic alterations in females, particularly in genes associated with synaptic functions, which closely resemble alterations observed in human brain regions vulnerable to AD, especially those related to memory processes. These transcriptomic changes correlate with NMDAR-mediated excitatory synapse weakening and to hippocampus-dependent memory deficits, particularly in females.

Previous work on AETA’s precursor CTF-η suggested that AETA might be increased in human AD patients because accumulation of CTF-η was reported in dystrophic neurites [51]. Most recently, a new study highlighted that an AD-linked mutation led to increase of AETA as well as Aβ production [21]. Here we show for the first time a significant accumulation of AETA in the hippocampus and PFC, both memory-linked regions of the brain. This increase seems to be due to post-transcriptional mechanisms since the AETA/APP protein ratio was increased, while APP protein and mRNA were not altered in AD brains. As AETA was just identified as a physiological modulator of NMDARs with strong consequences on NMDAR function [11], we would expect fine regulation of its availability with efficient clearance around synapses. Increased processing or reduced clearance of AETA could be at play in the AD brain.

Unfortunately, we cannot at present distinguish between the two possibilities due to several reasons. First, we see no possibility to monitor the activity of the different secretases participating to AETA production with the necessary resolution at hippocampal synapses. Also antibodies that specifically detect the β form of secreted AETA do not yet exist. Finally, there is currently no available information on the clearance mechanism of AETA in the brain. Interestingly, we recently reported that AETA production is increased upon enhanced neuronal activity [11]. As AD brains are known to exhibit hyperexcitability and even epileptic activity, especially during the early phase of the disease [42, 46], this could contribute to enhanced AETA levels. Since AETA was shown to dampen synaptic activity [11, 51], this activity-dependent increase in AETA levels might represent a physiologically-encoded control of network excitability that becomes over-solicited in the AD brain leading to abnormal synapse weakening. Why female AD brains exhibit more AETA than male AD brains has also to be further addressed, but a more complete understanding of this could help explain the prevalence of AD in women.

Our first analysis of the new AETA-m mouse model provides several important insights into the contribution of AETA in AD etiology. First, we did not observe any aggregation of AETA even following chronic over-expression. This could partly explain why this APP fragment was not easily detected in human AD brains remaining under the radar of neuropathologists for so long. Second, a neuron-centric analysis of RNA-seq data from AETA-m hippocampi, compared with human RNA-seq datasets, reveals that elevated AETA expression alters the hippocampal transcriptome, particularly affecting synaptic gene expression, resulting in a molecular profile that closely resembles that observed in human AD. As expected in a mouse model without overt neuroinflammation nor tau pathology, these molecular modifications were subtle but sufficient to provoke synapse dysfunction and could thus reflect early AD-like synaptopathy. Indeed, functionally, this translates to synapse weakening (less LTP, more LTD and spine loss), phenotypes also observed in human AD brains and in animal models of AD [40, 45]. Of note, in AD mouse models overexpressing APP, there will be *de facto* more AETA, as shown for the APPPS1 line [51], which might contribute to NMDAR-dependent synapse weakening observed in these models [25].

In our RNA-seq analysis, most genes that were down-regulated belong to pathways relevant for synapse function, consistent with our functional assessment of synapse weakening. Yet, we also observed a strong correlation of up-regulated genes when comparing the AETA-m and MSBB BM36 RNA-seq datasets. We could not find evidence of previous association of some of these up-regulated genes (FZD5, PLXNB2, CELSR1, FLNB) with AD pathology. However, there is previous evidence of the implication of the other genes (LEPR, ANO6, SGK3, FNBP1L, ATP11A and ADAM17) and their related signaling pathways in AD [6, 13, 14, 34, 36, 43, 50]. Although beyond the scope of the current study, analyzing further these upregulated genes, as well as the RNA-seq data beyond the neuronal signature, may yield additional insights into AETA-dependent brain perturbations. Fourth, the synaptic phenotypes observed in AETA-m mice upon chronic expression of AETA are similar to the acute action of AETA, most prominently affecting NMDAR signaling [11]. This latter observation provides key information. On one hand, it suggests that AETA remains within the scope of its primary physiological function at synapses, which is to weaken synapses via modulation of NMDAR ionotropic and ion flux-independent activities, although it is constantly available in brains of AETA-m mice. On the other hand, as we recently classified AETA as an endogenous modulator of NMDARs [11], it is likely to be dynamically regulated with specific production and clearance mechanisms. There is strong evidence that NMDAR dysfunction is a hallmark of AD [22, 24] and should be further explored as a therapeutic target in AD beyond the current FDA-approved treatment with the NMDAR antagonist memantine [15]. Once better identified, exploiting this endogenous regulation if AETA production and clearance or directly interfering with AETA’s action on NMDARs could represent innovative therapeutic strategies to normalize synapse function in AD patient brains.

Finally, while synapse function, spine number and P38 signaling were observed in both sexes, we observed a sexual dimorphism with several other markers of AD. Indeed, astrocyte and microglia number, the synaptic transcriptional signature and hippocampus-dependent memory retrieval were only significantly modified in female AETA-m mice. This latter deficit emerged when the delay of retention was increased (24h vs 7days) or when the task was more demanding (small pool vs larger pool). These memory deficits may reflect consolidation deficits reminiscent of prodromal AD [30, 47]. Surprisingly, while male AETA-m mice exhibited similar synapse weakening, they did not display significant memory deficits in these tasks. This could suggest that the memory deficits might also require astrocyte and/or microglial reactivity, only observed in females, to become evident. We currently do not hold an explanation for this sexual dimorphism in this mouse line. It does not seem to be due to different levels of human AETA expression as similar quantities were detected in both sexes, at least in the hippocampus. RNAseq data suggest that female hippocampi adapt the synaptic environment differently from male hippocampi to a chronic increase of AETA. Sex-specific memory deficits, prevailing in females, have also been observed in several mouse models of AD [5, 16, 32, 39, 52]. Explaining this sexual dimorphism, which currently remains enigmatic also in more classic AD mouse models, might help provide insights as to why AD is more prevalent in women.

Our study provides the first compelling evidence that AETA contributes to synaptic dysfunction in AD both at the transcriptional and functional levels, particularly modifying NMDAR signaling. Our findings align with and expand upon emerging views that synaptic failure in AD arises from multiple, potentially interacting molecular pathways beyond Aβ. Targeting AETA production, clearance or its interaction with NMDARs may represent a promising therapeutic strategy to restore synaptic function in prodromal AD [41], before irreversible neurodegeneration occurs.

## Author Contributions

JD performed immunoblotting, field and patch-clamp electrophysiology, behavior, mouse breeding and genotyping. CG performed immunoblotting, immunohistochemistry, Golgi Cox staining and analysis. SM, SMY and HM performed field electrophysiology. SM and HM performed patch camp experiments. AL, DB and LB performed qPCR on mouse tissue and tau immunoblotting. BA performed behavior, mouse breeding and genotyping. GP performed Golgi Cox staining and analysis, behavior and qPCR on human tissue. MT, KL, HL, HM and BM performed bioinformatics analysis of RNA-seq. PP and HM supervised and analyzed patch-clamp electrophysiology work. IB and HM supervised and analyzed behavior work. MW characterized AETA-m model and supervised human and mouse immunoblotting. HM piloted all aspects of study and manuscript writing with JD and contribution of other authors.

## Acknowledgments

We thank the Neuro-CEB Neuropathology network, which includes: Dr Franck Letournel (CHU Angers), Dr Marie-Laure Martin-Négrier (CHU Bordeaux), Dr Maxime Faisant (CHU Caen), Pr Catherine Godfraind (CHU Clermont-Ferrand), Pr Claude-Alain Maurage (CHU Lille), Dr Vincent Deramecourt (CHU Lille), Dr Mathilde Duchesne (CHU Limoges), Dr David Meyronnet (CHU Lyon), Dr Clémence Deltei (CHU Marseille), Pr Valérie Rigau (CHU Montpellier), Dr Fanny Vandenbos-Burel (Nice), Pr Danielle Seilhean (CHU PS, Paris), Dr Susana Boluda (CHU PS, Paris), Dr Isabelle Plu (CHU PS, Paris), Dr Dan Christian Chiforeanu (CHU Rennes), Dr Florent Marguet (CHU Rouen), Dr Béatrice Lannes (CHU Strasbourg), for providing the human PFC samples. We thank patient associations and research foundations that support the Neuro-CEB Biobank: ARSLA, CSC, France DFT, Fondation France SEP, Fondation Vaincre Alzheimer, France Parkinson, PSP France, for providing the human hippocampi samples. The authors thank Dr JGJ van Rooji, Erasmus University Medical Center Rotterdam, the Netherlands, for kindly providing additional information on NBB-HPC RNAseq database. The authors also thank the technical support of the UCA GenomiX. The authors acknowledge the flow cytometry or microscopy facility from the « Institut de Pharmacologie Moléculaire et Cellulaire » part of the « Microscopie Imagerie Côte d’Azur » GIS IBiSA labeled platform. We thank Dr Camilla Giudici, German Center for Neurodegenerative Diseases (DZNE) Munich, Germany, for help with AETA-m mouse characterization. We thank Julien Fassy, IPMC, for help with qPCR of human tissues.

