## Supplementary Figures for "AETA peptide contributes to Alzheimer’s disease signature of synapse dysfunction": R2_SI _Dunot.pdf

**Journal: Acta Neuropathologica**

Jade Dunot[1*], Carine Gandin[1], Marin Truchi[1], Giulia Pirro[1], Sébastien Moreno[1], Agathe Launay[2], Benjamin Azoulay[1], Hugo Landra[1,3], Sandy Ma Yishan[1,3], Luc Buée[2], Kevin Lebrigand[1], Paula A. Pousinha[1], David Blum[2], Bernard Mari[1], Ingrid Bethus[1], Michael Willem[4] and Hélène Marie[1*].

[1] Université Côte d'Azur, CNRS, INSERM, Institut de Pharmacologie Moléculaire et Cellulaire, Valbonne, France ; [2] Univ. Lille, INSERM, CHU Lille, UMR-S1172 LilNCog - Lille Neuroscience & Cognition, Lille, France; [3]Munich Cluster for Systems Neurology (SyNergy), Munich, Germany; [4] Biomedical Center (BMC), Division of Metabolic Biochemistry, Faculty of Medicine, Ludwig-Maximilians-Universität München, Munich, Germany

* Corresponding authors : Hélène Marie & Jade Dunot

Figures S1-S6

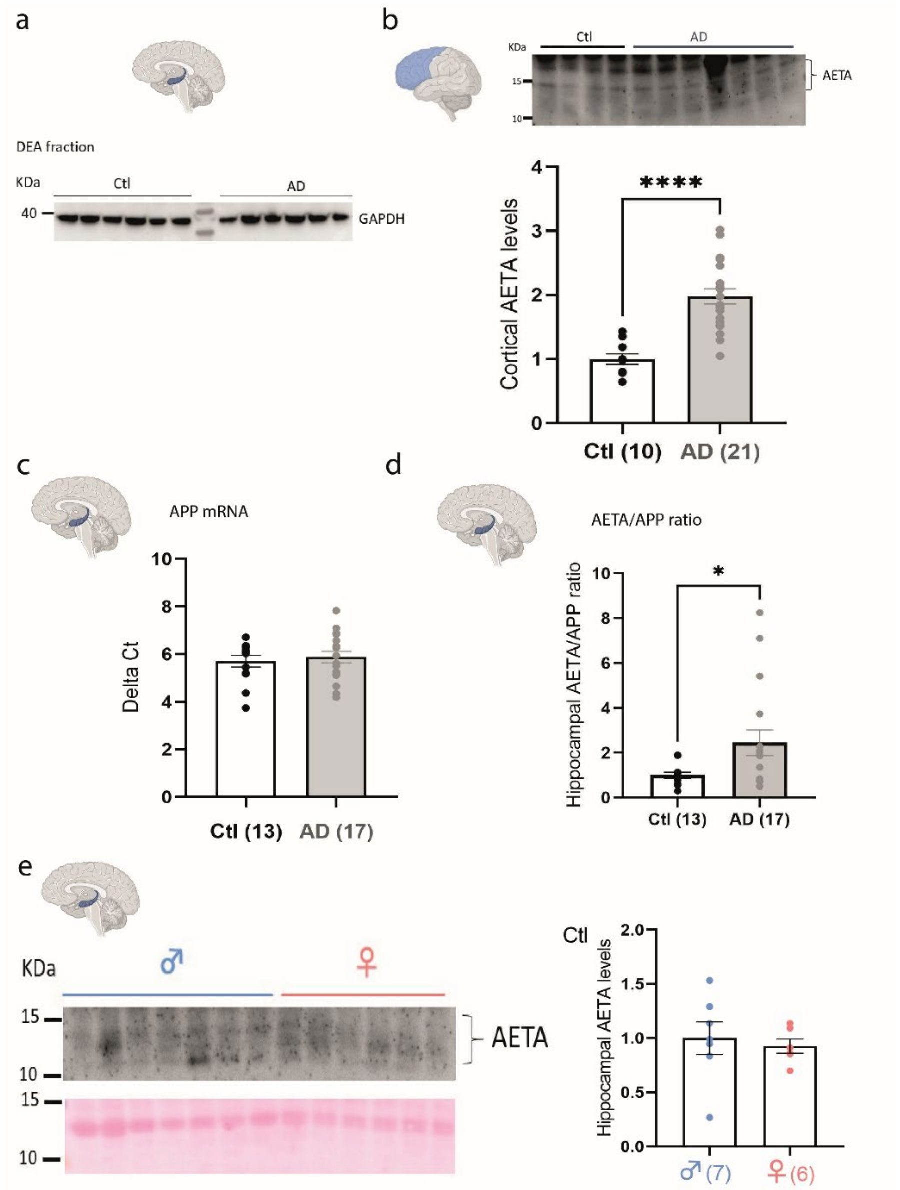


**Figure S1 (related to Figure 1): Additional AETA and APP quantification in hippocampi of AD patients and quantification of AETA in prefrontal cortex (PFC) of AD patients. (a)** General tissue integrity was confirmed by examples of GAPDH immunoblotting in DEA fractions of protein lysates. **(b)** Immunoblot (2D9 antibody) examples and quantification of AETA levels in PFC of AD patient vs control subjects (relative levels normalized to age-matched control (Ctl) subjects; normalized to actin loading control – see full blots used for analysis provided in Supplemental material). **(c)** Quantification of APP transcript by qPCR in hippocampi of human control subjects (Ctl) and AD brains. **(d)** The AETA/APP ratio was calculated from results obtained in graphs of Figure 1b-c. **(e)** Immunoblot (2D8 antibody and ponceau staining for normalization) examples (left) and quantification of AETA levels in hippocampi of male vs female control subjects (right). N= number of human samples; each sample was tested in triplicate and presented as mean. Error bars represent s.e.m.; * $p < 0.05$; **** $p < 0.0001$. Statistics: Welch's test for b and e, and Mann-Whitney test for c et d. See full statistics in Supplemental Table S9.

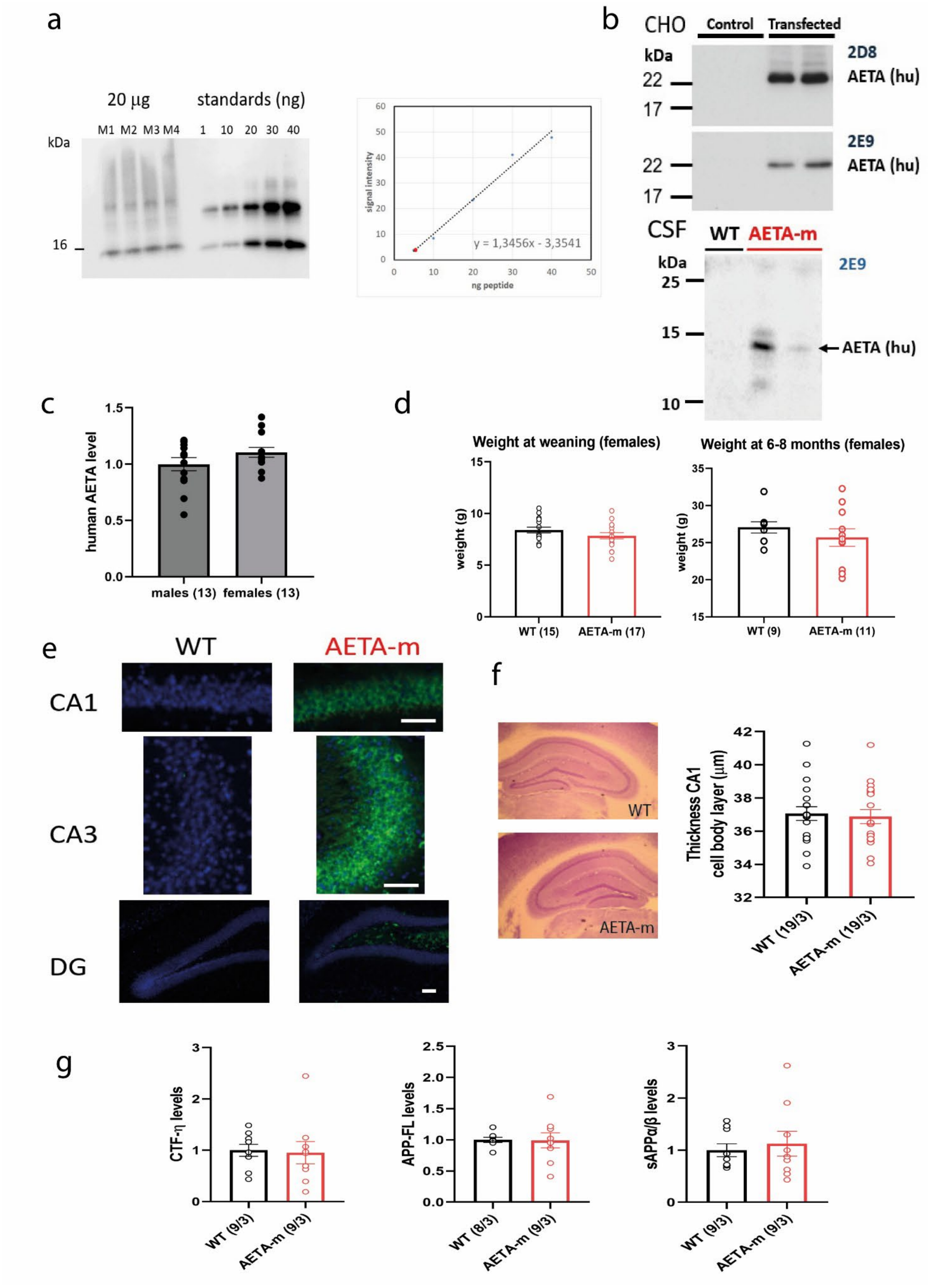
a
20 µg
standards (ng)
M1 M2 M3 M4 1 10 20 30 40
kDa
16
signal intensity
ng peptide
y = 1,3456x - 3,3541
b
CHO
Control
Transfected
kDa
2D8
AETA (hu)
2E9
AETA (hu)
CSF
WT
AETA-m
2E9
AETA (hu)
c
human AETA level
males (13)
females (13)
d
Weight at weaning (females)
Weight at 6-8 months (females)
weight (g)
WT (15)
AETA-m (17)
WT (9)
AETA-m (11)
e
WT
AETA-m
CA1
CA3
DG
f
WT
AETA-m
Thickness CA1 cell body layer (µm)
WT (19/3)
AETA-m (19/3)
g
CTF-η levels
WT (9/3)
AETA-m (9/3)
APP-FL levels
WT (8/3)
AETA-m (9/3)
sAPPα/β levels
WT (9/3)
AETA-m (9/3)

**Figure S2 (related to Figure 2): Additional characterization of AETA-m mice overexpressing secreted human AETA in the brain. (a)** (left) Immunoblot for quantification (2E9 antibody) of recombinant human AETA in hippocampi of four male AETA-m mice (6 months old; M1-M4). The quantification was made against a standard curve of synthetic AETA (1, 10, 20, 30, 40 ng). Note that recombinant AETA seems to partially dimerize both in brain and *in vitro*. (Right) Both bands of the synthetic peptide were used to generate the standard curve and calculate the average AETA quantity using the equation shown in graph (ng/$\mu$g protein; red dot on graph represents average quantity of human AETA calculated as 4,07 signal intensity in 20 $\mu$g of extracted protein; which gives 0,27 ng/$\mu$g using the provided equation and divided by 20 to obtain per quantity per $\mu$g). **(b)** Confirmation of secretion of recombinant human AETA: Detection of secreted human AETA-mycflag in medium of CHO transfected cells (AETA) and not in untransfected cells (control) by immunoblotting with 2D8 and 2E9 (top). Detection of  human AETA in CSF of female AETA-m mice and not of WT mice by immunoblotting with 2E9 (bottom).  **(c)** Quantification of human AETA levels (2E9 antibody on DEA lysate) in hippocampi of 6-8 months old female and male AETA-m mice. **(d)** Weight of female WT and AETA-m mice at weaning (21 days after birth) and at 6-8 months of age. **(e)** Representative examples of staining of human AETA peptide with 2E9 in CA1, CA3 cell body layers, and DG showing stronger expression within the CA layers in AETA-m mice compared to WT mice. . Nuclei were stained with DAPI (blue). Scale bars represent 50 $\mu$m. **(f)** Example of cresyl violet staining of hippocampi of male WT and AETA-m mice and measure of CA1 cell body layer thickness in these slices. **(g)** Quantification of endogenous levels of CTF-$\eta$, full-length APP (APP-FL) and sAPP $\alpha/\beta$ in WT and AETA-m mouse brains by immunoblotting. Full blots used for analysis are provided in Supplemental material. Three mouse brains per genotype were processed in triplicates.  n= mice for c-d; n/N= slices/ mice for f; n/N= replicates/ mice for g.  Error bars represent s.e.m. Statistics: Mann-Whitney test for c,d g left and e right. Student t-test for f and g center. See full statistics in Supplemental Table S11.

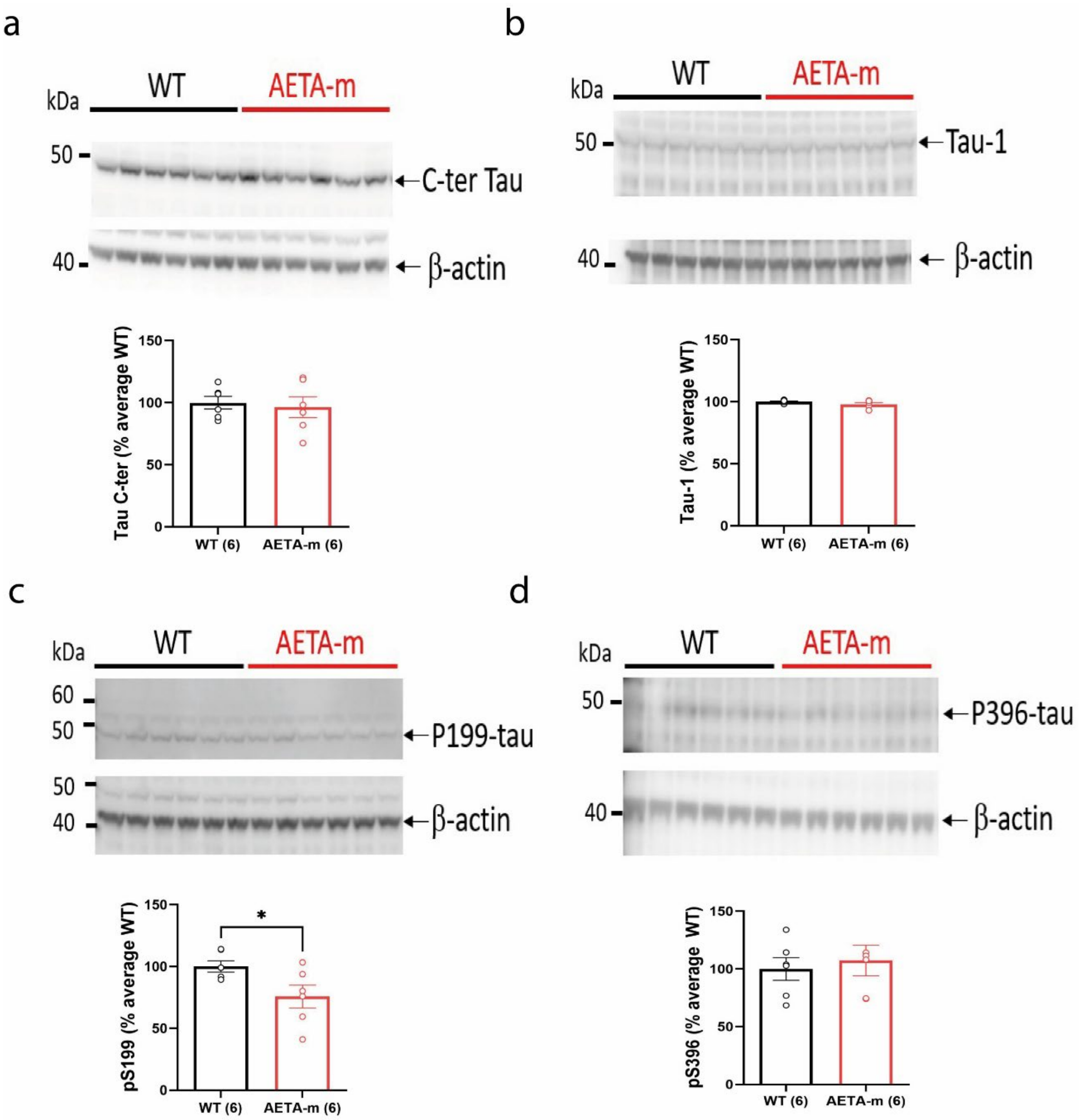


**Figure S3 -Tau levels and Tau phosphorylation do not increase in hippocampi of adult female AETA-m mice.** Immunoblots and associated quantification of endogenous levels of tau with C-terminal (a) and Tau-1 (b) antibodies and of endogenous tau phosphorylation at positions (c) S199 and (d) S396 in 6-8 months old female mice. Full blots used for analysis are provided in Supplemental material. n= mice. Error bars represent s.e.m.; * $p < 0.01$. Statistics: Student t-test. See full statistics in Supplemental Table S12.

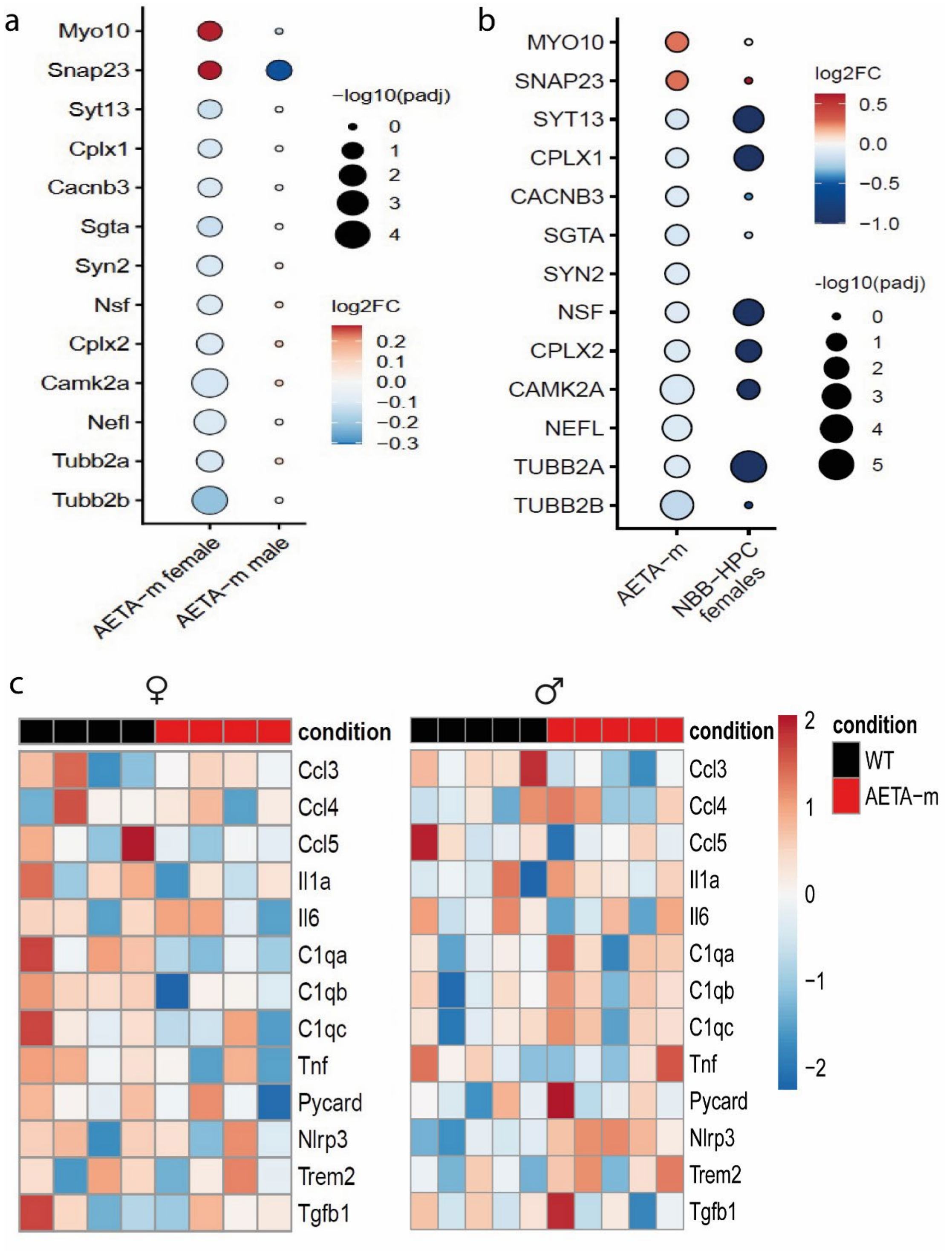


**Figure S4 (related to Figure 2 and 3) Additional analysis of bulk RNA sequencing and comparison to human NBB-HPC database. (a)** Lack of molecular signature of synaptic genes pertinent to AD in male AETA-m hippocampi. **(b)** Comparison of synaptic molecular signature between female AETA-m hippocampi and female-only NBB-HPC data. **(c)** Levels of mRNAs pertaining to neuroinflammation-related genes were unchanged in female (left) and male (right) hippocampi of AETA-m mice. For (c), see full statistics in Supplemental Tables S4 and S7.

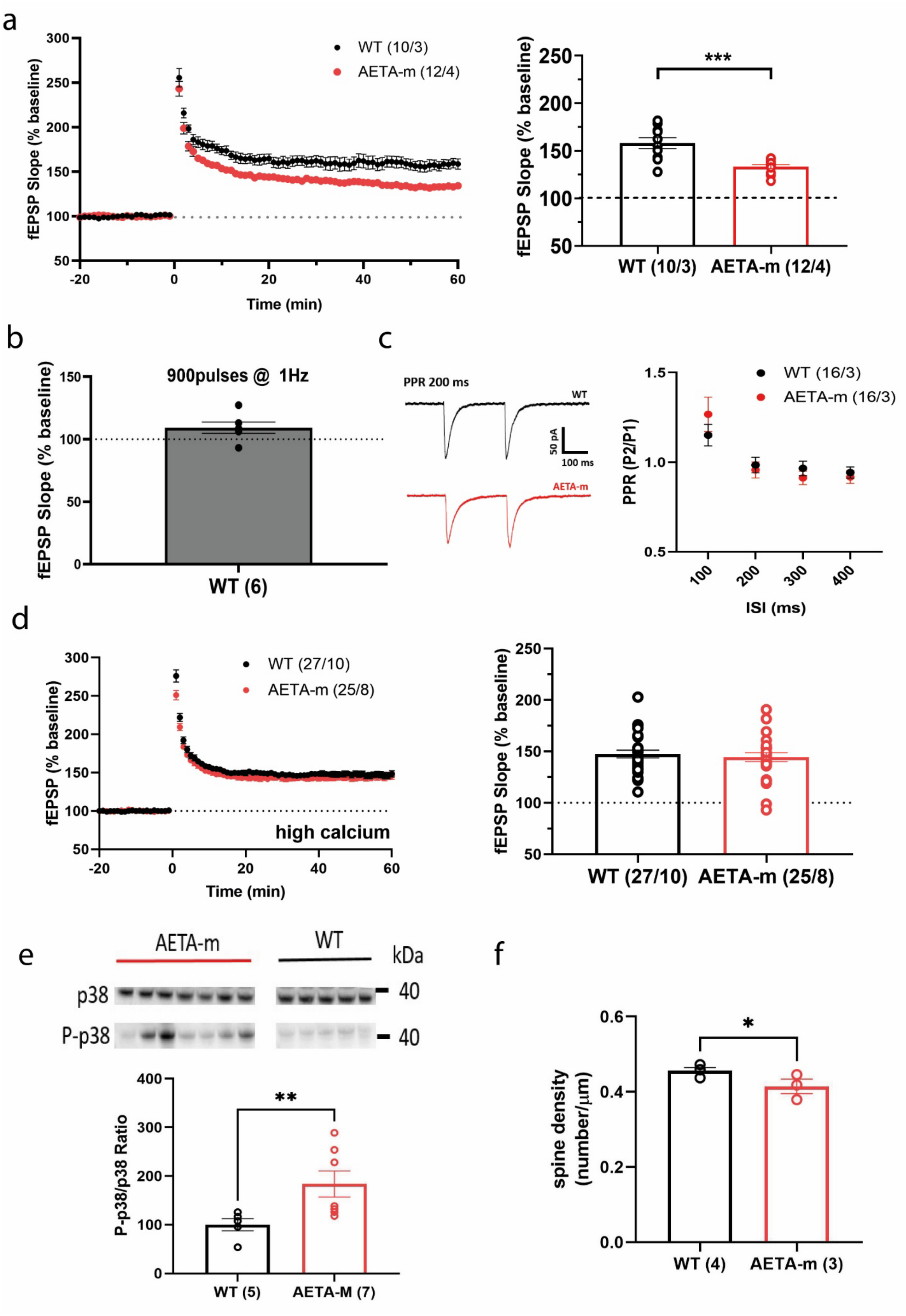
a
WT (10/3)
AETA-m (12/4)
fEPSP Slope (% baseline)
Time (min)
***
WT (10/3)
AETA-m (12/4)
b
900pulses @ 1Hz
WT (6)
c
PPR 200 ms
WT
AETA-m
50 pA
100 ms
WT (16/3)
AETA-m (16/3)
PPR (P2/P1)
ISI (ms)
d
WT (27/10)
AETA-m (25/8)
fEPSP (% baseline)
high calcium
Time (min)
fEPSP Slope (% baseline)
WT (27/10)
AETA-m (25/8)
e
AETA-m
WT
kDa
p38
P-p38
40
40
P-p38/p38 Ratio
**
WT (5)
AETA-M (7)
f
*
spine density (number/μm)
WT (4)
AETA-m (3)

**Figure S5 (related to Figure 4): Additional data of synapse function, plasticity and spine density in CA1 pyramidal neurons of AETA-m mice. (a)** Time course (left; LTP induction at time 0) and bar graph (right, 45-60 min post-induction) of long-term potentiation of fEPSP slope (% baseline) at CA3-CA1 synapse in hippocampal slices of male WT and AETA-m mice. **(b)** LTD could not be induced in adult female WT mice. **(c)** Representative traces (left) and paired pulse ratio (pulse 2/pulse 1) analysis (right) measured in CA1 pyramidal neurons of female WT and AETA-m mice. **(d)** Time course (left, LTP induction at time 0) and bar graph (right, 45-60 min post-induction) of long-term potentiation of fEPSP slope (% baseline) at CA3-CA1 synapse in hippocampal slices of male WT and AETA-m mice recorded with aCSF containing high calcium. **(e)** Example immunoblots of p38 and phosphorylated p38 (P-p38) and P-p38/p38 ratio quantification in male WT and AETA-m mice. Samples were tested in quadruplicate. **(f)** Spine density measured on dendrites of the stratum radiatum of CA1 pyramidal neurons of male WT and AETA-m mice. Error bars represent s.e.m.; * $p < 0.05$; ** $p < 0.01$; *** $p < 0.001$. n/N= slices/mice for a and d; n/N= cells/mice for c; n= number of mice for e-f. Statistics : Student t-test for a and d; Two-way ANOVA for c; Mann-Whitney for e and f. See full statistics in Supplemental Table S14.

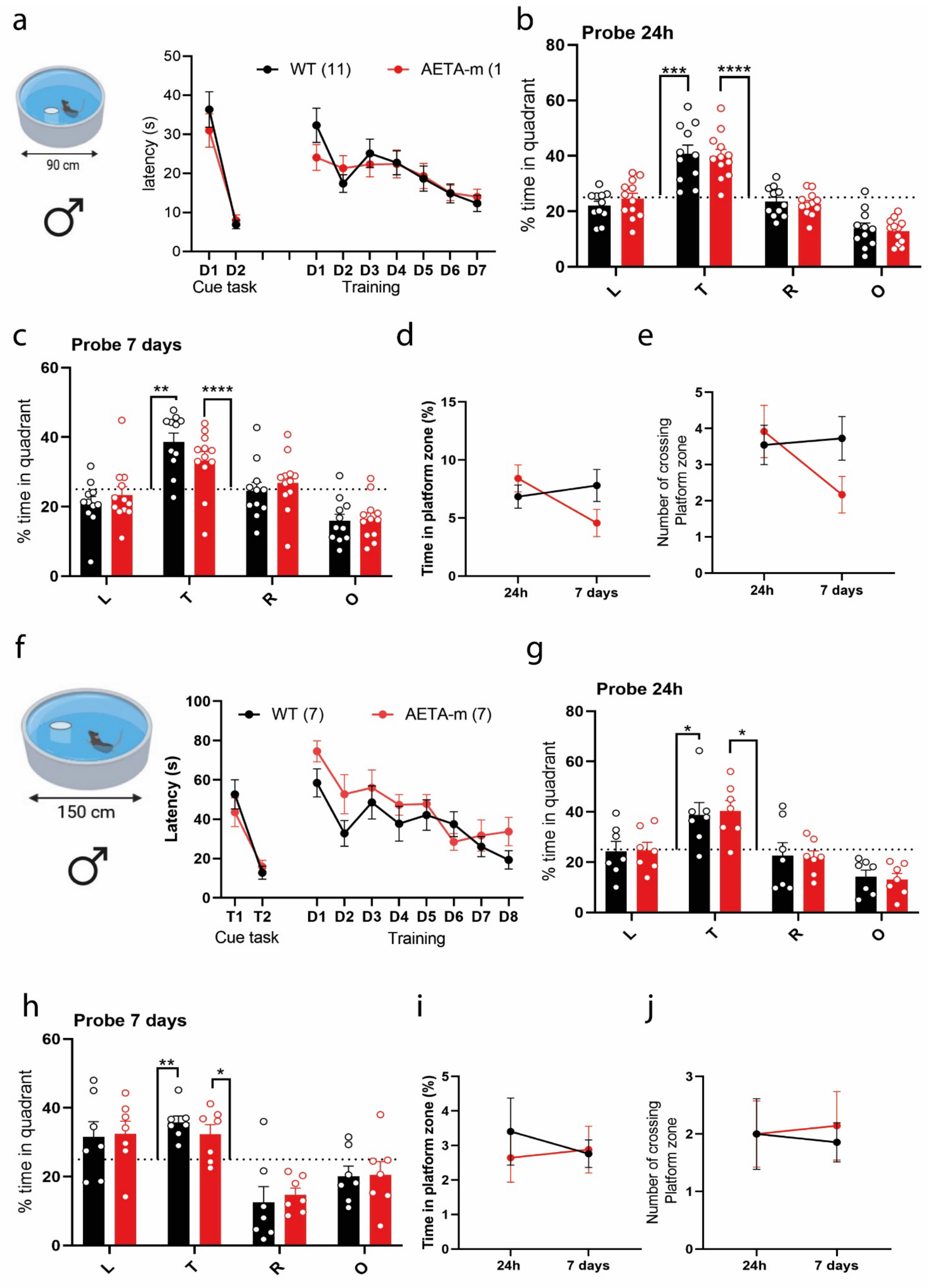
a
90 cm
latency (s)
WT (11)
AETA-m (1
D1 D2
Cue task
D1 D2 D3 D4 D5 D6 D7
Training
b
Probe 24h
% time in quadrant
L T R O
c
Probe 7 days
% time in quadrant
L T R O
d
Time in platform zone (%)
24h 7 days
e
Number of crossing Platform zone
24h 7 days
f
150 cm
Latency (s)
WT (7)
AETA-m (7)
T1 T2
Cue task
D1 D2 D3 D4 D5 D6 D7 D8
Training
g
Probe 24h
% time in quadrant
L T R O
h
Probe 7 days
% time in quadrant
L T R O
i
Time in platform zone (%)
24h 7 days
j
Number of crossing Platform zone
24h 7 days

**Figure S6 (related to Figure 5): Long-term memory in male AETA-m mice. (a)** Latency (s) during cue task and training in 90 cm Morris water maze pool. **(b-c)** % time spend in quadrants (T: target, L: left, R: right, O: opposite) during probe test (b) at 24h, (c) at 7 days. **(d)** Time spent in platform zone at 24h and 7 days. **(e)** Number of platform crossing at 24h and 7 days. **(f)** Latency (s) during cue task and training in 150 cm Morris water maze pool. **(g-h)** % time spend in quadrants during probe test at 24h and at 7 days. **(i)** Time spent in platform zone at 24h and 7 days. **(j)** Number of platform crossing at 24h and 7 days. N= mice. Error bars represent s.e.m.; *$p < 0.01$; ** $p < 0.001$; *** $p < 0.0001$; **** $p < 0.0001$; against chance level (25%, dotted line on graph). Statistics: two-way ANOVA. See full statistics in Supplemental Table S16.
